# The HBV basal core promoter mutation confers a replicative advantage and transcriptionally reprograms hepatocytes toward HCC subtypes

**DOI:** 10.64898/2026.09.22.752323

**Authors:** Leon L. Seifert, Georgios Dangas, Hoyin Chu, Yingpu Yu, Kosuke Ogata, Xupeng Hong, Mengyin Zhang, Yichen Zhou, Chenhui Zou, Alireza Ramandi, Corrine Quirk, Evgenia Moschogianni, Catherine A. Freije, Antonis Athanasiadis, Leonardo Gonzales, Luis Chiriboga, Clifton Fulmer, Hong Hur, Manoj Kandpal, Anna S. Lok, Caleb Lareau, William M. Schneider, Charles M. Rice, Eleftherios Michailidis, Ype P. de Jong

**Affiliations:** Laboratory of Virology and Infectious Disease, The Rockefeller University, New York, NY, USA; Center for Clinical and Translational Science, The Rockefeller University Hospital, New York, NY, USA; Laboratory of Biochemical Pharmacology, Department of Pediatrics, Emory University, Atlanta, GA, USA; Computational and Systems Biology Program, Memorial Sloan Kettering Cancer Center, New York, NY, USA; Department of Molecular Systems BioAnalysis, Kyoto University, Kyoto, Japan; Division of Gastroenterology and Hepatology, Weill Cornell Medicine, New York, NY, USA; Department of Pathology, NYU School of Medicine, New York, NY, USA; Department of Pathology and Laboratory Medicine, Cleveland Clinic, Cleveland, Ohio, USA; Department of Research Bioinformatics, Center for Clinical and Translational Science, The Rockefeller University Hospital, New York, NY, USA; Department of Internal Medicine, Michigan Medicine, University of Michigan, Ann Arbor, Michigan

**Author notes:** Authors contributed equally (Charles M. Rice, Eleftherios Michailidis, Ype P. de Jong). **Correspondence:** L.L.S. and C.M.R.: The Rockefeller University, 1230 York Avenue, New York, NY 10065. E.M.: Emory University, Health Sciences Research Building II, 1750 Haygood Drive NE, Atlanta, GA 30322. Y.P.J.: Division of Gastroenterology and Hepatology, Weill Cornell Medicine, 413 East 69th Street, New York, 10021 NY.

**Keywords:** hepatitis B virus, viral hepatitis, hepatocellular carcinoma

## Abstract

**Background:** Hepatitis B virus (HBV) basal core promoter (BCP) and precore (PC) mutations occur during chronic HBV infection with the BCP-mutant being associated with increased hepatocellular carcinoma (HCC) risk.

**Objectives:** The effects of these mutants on viral replication, hepatocyte biology, and carcinogenesis remain poorly defined. To address this, we characterized BCP- and PC-mutants using human hepatocyte chimeric mice and asked whether the resulting infection-induced transcriptomes correspond to subsets of human HBV-related hepatocellular carcinomas (HBV-HCCs).

**Design:** Isogenic wild-type (WT), BCP- and PC-mutants of HBV genotypes A, C, and D (HBV-A, -C, -D) were generated from recombinant covalently-closed circular DNA to infect chimeric mice. HBV-HCC transcriptomic datasets from The Cancer Genome Atlas (TCGA) were used to stratify association of HBV variants with HCC subtypes.

**Results:** WT, BCP-mutant, and PC-mutant HBV sequences remained genetically stable. The BCP mutation, but not the PC mutation, accelerated the rise in serum viremia in HBV-D- and HBV-A-infected chimeras; in HBV-C, acceleration required both (PC+BCP) mutations. Further comparisons of HBV-D variants revealed that the BCP-mutant increased intrahepatic viral DNA, viral protein expression, and upregulated cancer-related pathways, including transcripts associated with a subset of HBV-HCCs. Analysis of HBV-HCC samples from the TCGA revealed that tumors often harbor a mixture of WT and mutant transcripts, and that WT- and BCP-mutant-associated HCCs exhibit distinct transcriptional profiles.

**Conclusions:** The HBV BCP-mutant directly perturbs hepatocyte homeostasis via virus-intrinsic mechanisms, selectively activating cancer-related pathways and defining a molecularly distinct subset of HBV-HCC. These findings suggest that HBV variants form distinct subcategories of HBV-HCCs.

## Introduction

Worldwide chronic hepatitis B virus (HBV) infection causes approximately 1.1 million deaths annually, mostly from liver cirrhosis and hepatocellular carcinoma (HCC).[1] Chronic HBV infection is divided into clinical phases defined by the presence or absence of the HBV e-antigen (HBeAg). Initial phases are HBeAg-positive, while later phases are HBeAg-negative, marked by the loss of HBeAg and seroconversion.[2] In HBeAg-negative patients, the most common enriched mutations are the precore mutation (PC-mutant, G1896A) and two basal-core promoter mutations that occur in combination (BCP-mutant, A1762T/G1764A).[3] The PC-mutant abolishes HBeAg production by introducing a stop codon in the HBV precore messenger RNA (pcRNA), whereas BCP mutations decrease HBeAg levels by about 50%.[4–8] The BCP-mutant, which is linked to increased risk of HCC development, has shown enhanced viral replication in cell culture systems.[9–15]

As HBV infected individuals can maintain normal liver function for decades, HBV is considered non-cytopathic.[16–19] The prevailing assumption is that liver injury and carcinogenesis are driven by adaptive antiviral immune responses rather than direct viral effects.[20–25] However, HBV may have direct cytopathic effects upon enhanced virus replication and increased protein production.[26] Some experimental studies have challenged the notion that HBV remains non-cytopathic in the presence of the BCP or PC-mutants.[27–29]

Given HBV’s strict human tropism, only mice engrafted with primary human hepatocytes (PHHs) support HBV infection.[30,31] Previous HBV-studies have focused on patient isolates[27–31], which exhibit substantial genetic variability between and within individual patients[32,33], precluding controlled comparisons. Cell culture-derived stocks can provide genetically defined viral clones but generating stable cell lines remains laborious and inefficient.[34]

In this study, we aimed to compare how wild-type (WT) and isogenic BCP- or PC-mutant infections affected PHH engrafted in immunodeficient *Fah^−/−^*NOD *Rag1^−/−^ Il2rg^null^* (huFNRG) hepatocyte chimeric mice.[35,36] As clinical outcomes differ by HBV genotype[37], we produced BCP- and PC-mutants from parental sequences of HBV genotypes A, C, and D. These were launched via intrahepatic injection (IHI) of recombinant covalently-closed circular DNA (rcccDNA) enabling defined infection of huFNRG mice with isogenic stocks. This allowed dissection of mutant-specific effects without the confounding influence of adaptive immune responses or heterogeneous HBV inocula. Proteomic and transcriptomic analyses showed upregulation of cancer-associated pathways in BCP-infected human hepatocytes, prompting a comparison with publicly available HCC datasets.

## Results

### Intrahepatic injection of recombinant HBV cccDNA yields genetically stable virus stocks, enabling isogenic infection comparisons

To generate infectious HBV harboring BCP and PC mutations, we employed a newly developed rcccDNA[38] intrahepatic injection (IHI) protocol in huFNRG mice. BCP and PC mutations were engineered into two prototype parental WT-sequences (HBV-A[39] and HBV-D[40]) and one published HBV-C sequence[41]. Following purification, rcccDNA was administered via IHI into huFNRG mice (**Fig. 1A**) and serum viremia was monitored with sequence-specific primers (**Table S1**). This yielded effective virus production and spread across HBV genotypes and mutants in 30/34 mice; four mice died before stable viremia was achieved (**Fig. 1B** and **Fig**. **S1**).

**Figure 1:**
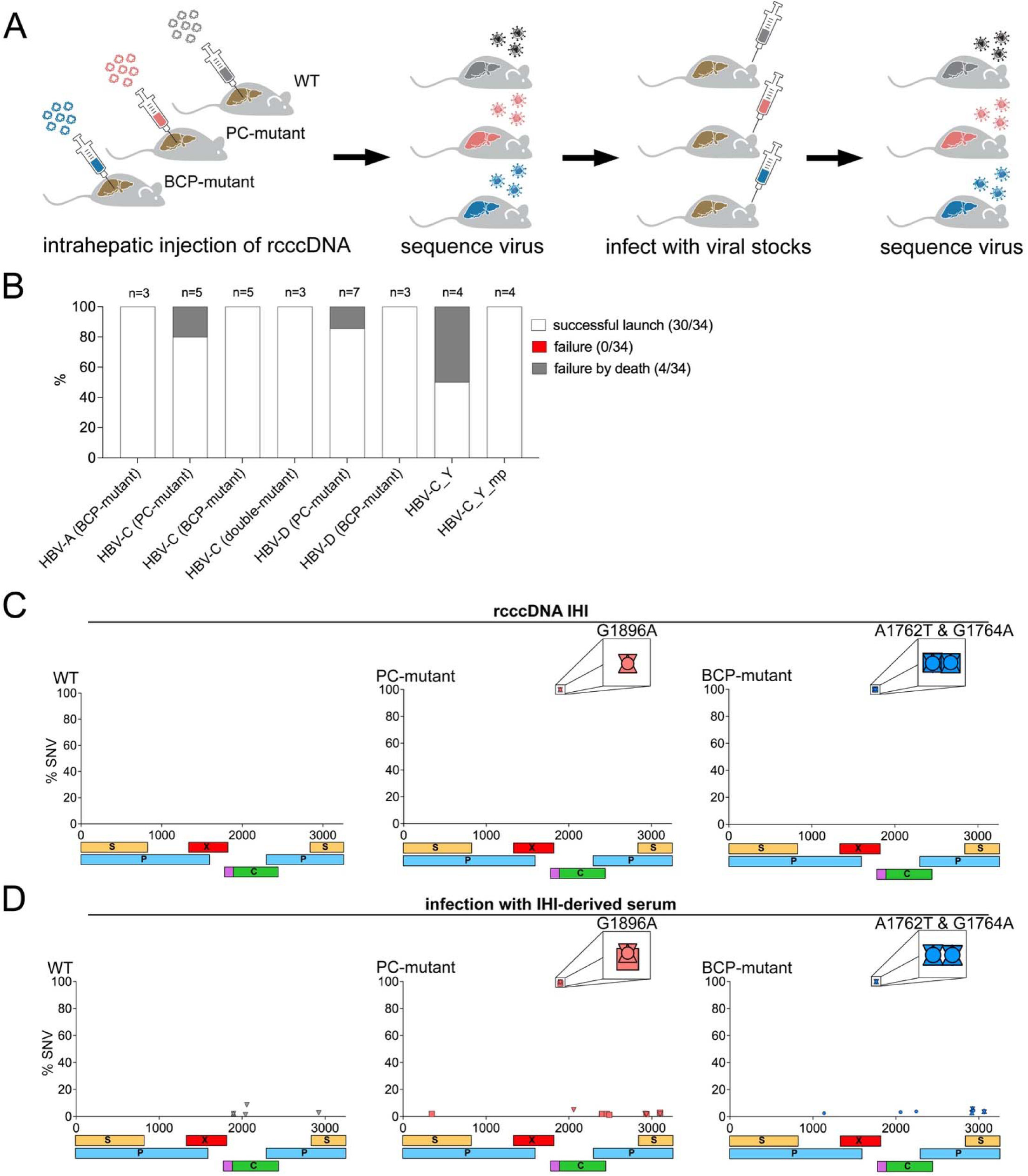
PC and BCP mutations do not revert to wild-type in immunodeficient mice. **(A)** Schematic overview of the workflow for generating isogenic infectious HBV stocks via IHI rcccDNA launch in huFNRG mice. Mice developed HBV viremia following IHI, and serum was used for sequencing analyses and as inoculum in downstream infection experiments. **(B)** Summary of HBV launches of isogenic HBV mutants via rcccDNA IHI. A “successful launch” was defined as serum viremia ≥ 10^5^ GE/mL with two consecutive increases in viral load. A “failure” was defined as the absence of a successful launch within 10 weeks after IHI. A “failure by death” indicates animals that died prior to week 10 post IHI without achieving a “successful launch”. **(C)** Sequencing analysis of HBV DNA isolated from serum of huFNRG mice following HBV rcccDNA IHI (HBV-C). Each data point represents an SNV, with each mouse represented by a different symbol (n= 3-4). The x-axis represents the position on the HBV genome, with the EcoRI restriction site in the S open reading frame (ORF) serving as the reference starting point. S = S-antigen ORF, P = polymerase ORF, X = X-protein ORF, C = core ORF (the pink extension represents the ORF unique to the precore protein). **(D)** Sequencing analysis of HBV DNA from serum of huFNRG mice infected with stocks generated in (C). Each data point represents an SNV, with each mouse represented by a different symbol (n= 3-4). *huFNRG mice = humanized Fah^−/−^ NOD Rag1^−/−^ Il2rg^null^ mice, IHI = intrahepatic injection, rcccDNA = recombinant covalently closed circular DNA, SNV = single nucleotide variation*.

We then tested whether the WT parental sequence would develop BCP or PC mutations or, conversely, whether such mutants would revert to WT in this *in vivo* model lacking adaptive immune pressure. At peak viremia, near-full-length amplicons were generated using genotype-specific primers (**Table S2**) and sequenced. The engineered BCP and PC mutations were detected in >99.8% of reads in mutant infected animals, with no *de novo* mutations seen with WT parental sequences (**Fig. 1C**). We also examined a patient-derived HBV-C[42–44] harboring both BCP and PC mutations. Consensus sequences were generated from the patient serum before (HBV-C_Y) and after passage in huFNRG mice (HBV-C_Y_mp). Both sequences showed no reversion WT after IHI (**Fig. S1** and **Fig. S2A**).

Finally, we examined whether infection of naïve huFNRG mice with serum from IHI-treated huFNRG mice would result in genetic adaptation of the virus. For these infection experiments, we used only highly humanized huFNRG mice (∼90% human hepatocytes, corresponding to ∼10 mg/ml serum human albumin[35]). Upon reaching peak viremia, the engineered BCP and PC mutations remained present in >98.5% of reads (**Fig. 1D** and **Fig. S2B** and **S2C**). Non-convergent single-nucleotide variants (SNVs) were detected at low frequency (<10%) in some animals, likely due to the presence of random variants in the bottlenecked low dose inocula which were then amplified upon passaging and expansion.

These findings demonstrate that BCP and PC mutants and their WT parental backgrounds are genetically stable in huFNRG mice, setting the stage for robust phenotypic comparisons.

### Effects of the BCP-mutation on the kinetics of serum viremia

We next infected huFNRG mice with a low viral inoculum (10^5^ GE). Viral kinetics differed, with WT HBV-A and -D viremia peaking around 10^9^ GE/ml and WT HBV-C peaking between 10^7^ and 10^8^ GE/ml. The HBV-A BCP-mutant exhibited accelerated serum viremia, reaching a higher peak than HBV-A WT and its PC-mutant, an effect even more pronounced for HBV-D (**Fig. 2A** and **Fig. S3A,C**). This accelerated increase in viremia of the HBV-D BCP-mutant was confirmed in three follow-up experiments, including huFNRG mice humanized with a different hepatocyte donor (**Fig. S4**). In contrast, HBV-C serum viremia levels were similar across WT, BCP- and PC-mutants (**Fig. 2A** and **Fig. S3B**). Given a frequent co-occurrence of BCP and PC mutations in HBV-C patients[3], we generated a “double-mutant” (DM: BCP+PC mutations) which showed an accelerated increase in serum viremia compared to WT HBV-C (**Fig. S5**).

**Figure 2:**
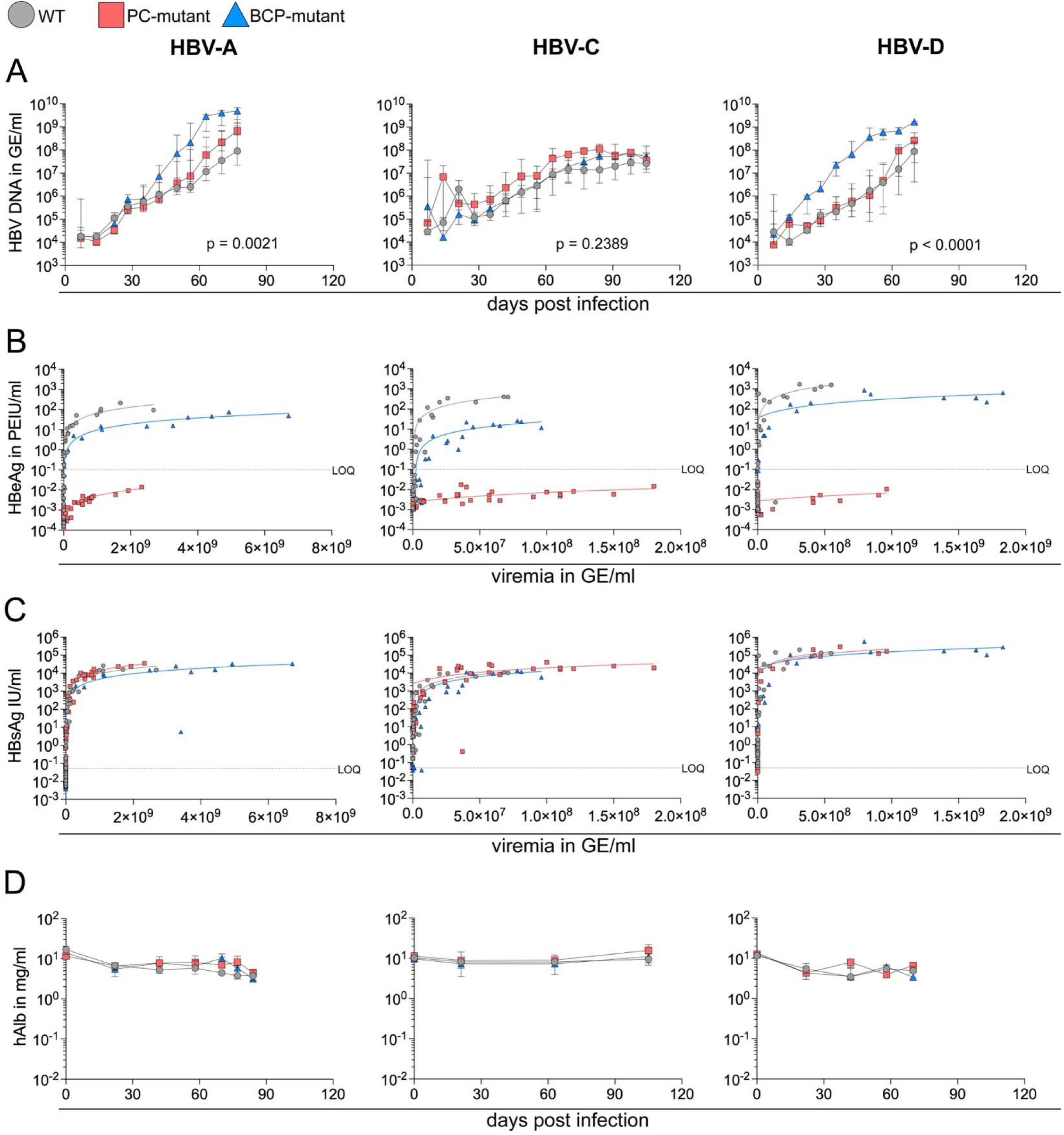
The BCP mutation confers a replicative advantage, varying by strain. Infection experiments with WT, BCP-mutant, or PC-mutant HBV across isolates of genotypes A, C, and D (n = 3-5 mice per group) in highly humanized huFNRG mice (serum human albumin ∼10 mg/ml at infection). Mice were infected with 10^5^ GE/mouse via retroorbital injection. **(A)** Serum HBV viremia over time. Data are shown as the median per group with the interquartile range. Statistical analysis was performed using a linear mixed-effects model fitted by restricted maximum likelihood (REML), including fixed effects for *group*, *time*, and *group × time* interaction and a random effect for subject. HBV-A (*time*: p = 0.0009; *group*: p = 0.0001; ***time x group*: p = 0.0021),** HBV-C (*time*: p = 0.0568; *group*: p < 0.0001; ***time x group*: p = 0.2839**), HBV-D (*time*: p < 0.0001; *group*: p < 0.0001; ***time x group*: p < 0.0001**). **(B)** Serum HBeAg levels, measured by chemiluminescence immunoassay, plotted against serum HBV viremia. Each symbol represents a single time point and animal. **(C)** Serum HBsAg levels, measured by chemiluminescence immunoassay, plotted against serum HBV viremia. Each symbol represents a single time point and animal. **(D)** Serum human albumin levels over time, measured via enzyme-linked immunosorbent assay. Data are displayed as median per group with interquartile range. *GE = genome equivalent, huFNRG mice = humanized Fah^−/−^ NOD Rag1^−/−^ Il2rg^null^ mice, IU = international units, LOQ = lower limit of quantification, PEIU = Paul-Ehrlich-Institute units*.

We then assessed the impact of the BCP-mutant on HBeAg levels. Normalizing to viremia levels for each individual mouse, HBeAg levels for BCP-mutants were reduced by 84.9% to 97% at peak viremia (**Fig. 2B, Fig. S6A-C,** and **Table S3**). As expected, HBeAg levels were below or at the limit of detection in PC-mutant infections. HBsAg levels were similar across WT, BCP- and PC-mutants (**Fig. 2C, Fig. S6D-F,** and **Table S4**).

Serum human albumin was used to monitor human graft stability (**Fig. 2D**). Importantly, no consistent differences were observed between WT and mutant infections. Serum human albumin levels still reflected the degree of liver humanization after HBV infection as assessed by fumarylacetoacetate hydrolase (FAH) staining (**Fig. S7**).

These data demonstrate that the BCP mutation alone accelerated serum viremia for HBV-A and HBV-D, whereas for HBV-C only the BCP+PC DM had this effect. HBeAg serum levels of BCP mutants were reduced by ∼90%, greater than previously described[6,8]; with levels undetectable for the PC mutants.

### Effects of HBV WT infection on human graft histology and transcription

We then examined the impact of HBV WT infections on the human hepatocyte graft, which was collected when mouse viremia had plateaued and almost all human hepatocytes were infected.[35,45–47] Histological examination of humanized areas (smaller, paler hepatocytes by H&E staining, **Fig. S8A**) revealed no significant differences in humanization between WT and uninfected livers (**Fig. S8B**). There was a slight increase in graft steatosis in HBV-C WT infection, but no significant WT differences in fibrosis (**Fig. S8C, S8D**).

We performed RNA-seq on bulk liver samples targeting 50 million reads per sample to account for human-murine chimerism. On average, 16% of reads could not be mapped and were discarded, resulting in 63% uniquely mapped human reads and 37% murine reads (**Fig. 3A**), with some animal-to-animal variability (**Fig. S9**). Human gene expression in WT HBV-A, HBV-C, and HBV-D infected hepatocytes clustered separately, demonstrating an HBV sequence-specific effect on the liver transcriptome (**Fig. 3B**). HBV WT infection clustered away from uninfected control livers, indicating that WT HBV infection alters the hepatocyte transcriptome (**Fig 3C**). Expression of some of the dysregulated genes (DEGs) in HBV infected huFNRG mice has previously been described to be altered in patients with HBV infection. For example, *ID1* has been shown to be downregulated by HBV[48] and expression was consistently lower in WT HBV-infected huFNRG mice.

**Figure 3:**
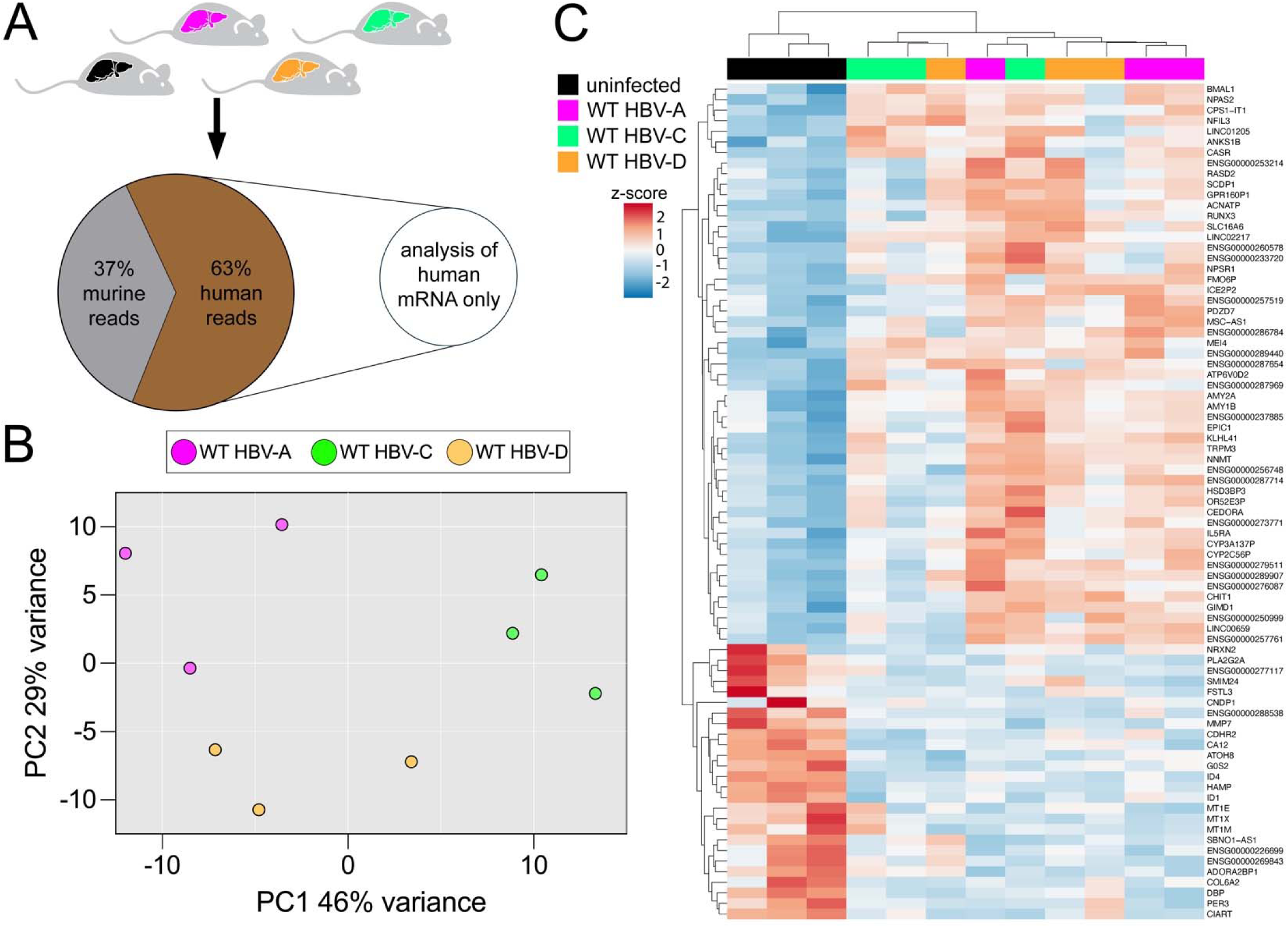
HBV WT infections cause limited human graft perturbations. **(A)** Livers of HBV-infected huFNRG mice were collected at peak plateau viremia (or corresponding time points in uninfected controls), total RNA was extracted, and RNA sequencing was performed. Shown are the relative proportions of reads mapping to the human (63%) or murine (37%) genome of all reads uniquely mapped to either reference genome. Only reads uniquely mapping to the human genome were used for downstream transcriptomic analyses. **(B)** Principal component analysis of mice infected with WT HBV-A, WT HBV-C, or WT HBV-D, based on the 1,000 most variable genes between uninfected and WT HBV infected samples. **(C)** Heatmap depicting differential human gene expression in livers from huFNRG mice infected with WT HBV-A, WT HBV-C, or WT HBV-D compared with uninfected controls (p-adj < 0.05, log2fc > |1.5|). *huFNRG mice = humanized Fah^−/−^ NOD Rag1^−/−^ Il2rg^null^ mice.*

These findings show that WT HBV infection did not cause histological changes but did alter expression of a limited number of genes, some of which were HBV WT genotype-dependent.

### Effects of the BCP-mutation on intrahepatic viral replication and antigen production

We decided to focus on HBV-D, where the BCP-mutant showed the most striking increase in serum viremia. Liver humanization, human graft steatosis, and hepatic fibrosis exhibited minor differences across HBV-D WT, BCP-, and PC-mutant infected mice (**Fig. 4A-C, Fig. S10A,B**). These minor histological differences contrasted with viral antigen staining, where BCP-mutant infected livers had higher expression of both HBsAg and HBcAg compared to WT and the PC-mutant (**Fig. 4A,D,E**).

**Figure 4:**
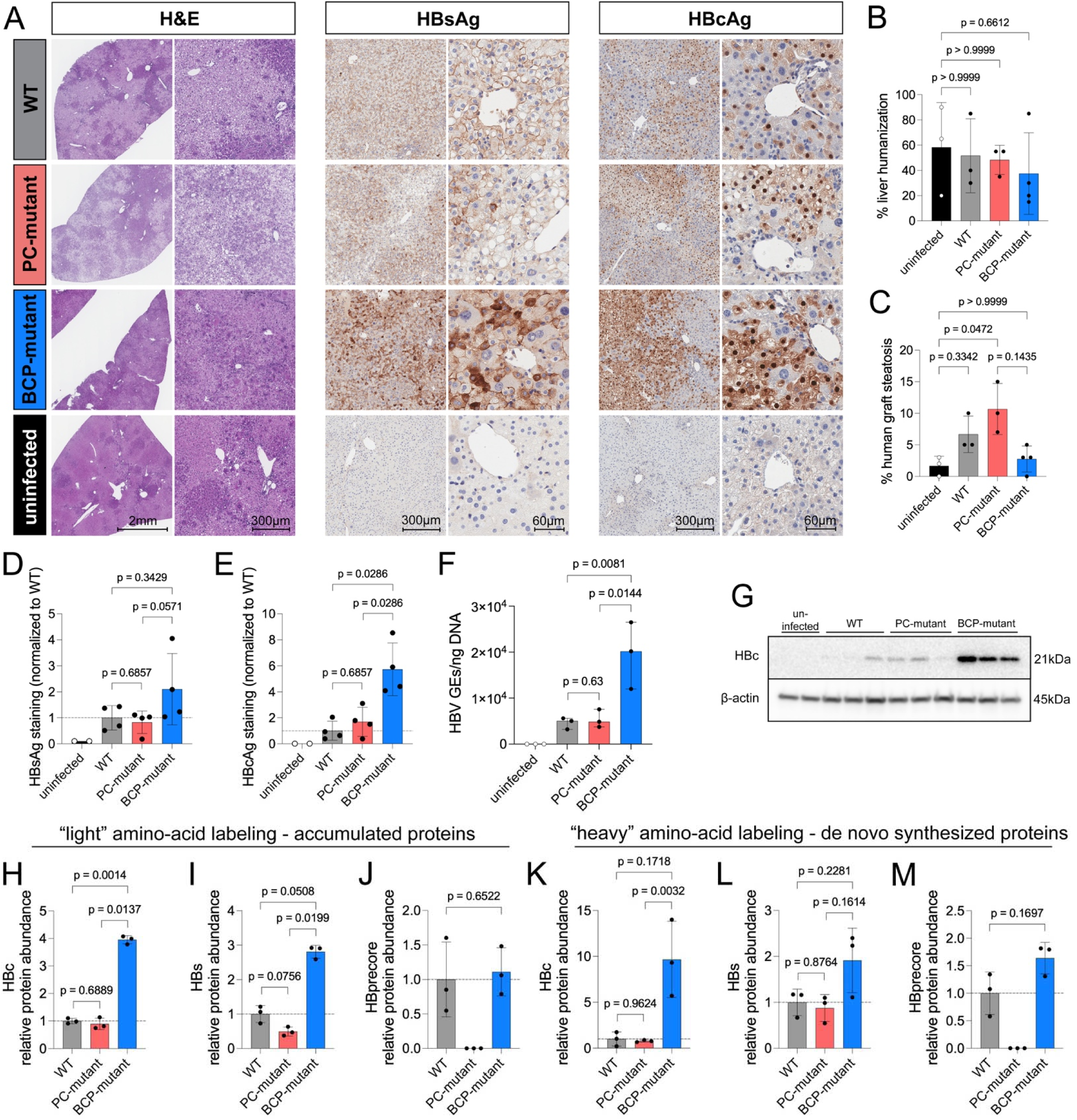
HBV-D BCP-mutant infection reshapes the human hepatocyte graft. **(A)** Representative hematoxylin and eosin (H&E), hepatitis B surface antigen (HBsAg), and hepatitis B core antigen (HBcAg) staining of liver sections from huFNRG mice infected with HBV-D WT, BCP-mutant, PC-mutant, and uninfected controls. **(B)** Percentage of human hepatocytes among all hepatocytes. Statistics: Kruskal-Wallis test with Dunn’s multiple comparisons test. Additional comparisons: WT vs. PC-mutant: p > 0.9999, WT vs. BCP-mutant: p > 0.9999, PC-mutant vs BCP-mutant: p > 0.9999. **(C)** Quantification of liver steatosis within the human hepatocyte graft. Statistics: Kruskal-Wallis test with Dunn’s multiple comparisons test. Additional comparisons: WT vs. PC-mutant: p > 0.9999, WT vs. BCP-mutant: p = 0.8607. **(D)** Quantification of HBsAg-positive area in the liver, normalized to the degree of humanization and WT. Statistics: two-tailed Mann-Whitney test. **(E)** Quantification of HBcAg-positive area in the liver, normalized to the degree of humanization and WT. Statistics: two-tailed Mann-Whitney test. **(F)** HBV DNA copy number as genome equivalents (GE) per ng of DNA in isolated mpPHHs. Statistics: pairwise comparisons via two-tailed lognormal Welch’s t-test after log10 transformation. **(G)** Western blot of HBcAg expression in non-cultured, freshly isolated PHHs. **(H-J)** Relative protein abundance of HBV core protein **(H)**, preS1 protein **(I)**, and precore protein **(J)** in mpPHHs infected with HBV-D, normalized to WT infection (“light amino acid labeled” peptides). Statistics: lognormal Brown-Forsythe and Welch ANOVA tests with Tukey test for multiple comparisons after log10 transformation **(H** and **I)**, pairwise comparison (WT vs. BCP mutant) via two-tailed lognormal Welch’s t test after log10 transformation **(J)**. **(K-M)** Relative protein abundance of HBV core protein **(K)**, preS1 protein **(L)**, and precore protein **(M)** in mpPHHs infected with HBV-D, normalized to WT infection (“heavy amino acid labeled” peptides). Statistics: lognormal Brown-Forsythe and Welch ANOVA tests with Tukey test for multiple comparisons after log10 transformation **(L and L)**, pairwise comparison (WT vs. BCP mutant) via two-tailed lognormal Welch’s t test after log10 transformation **(M)**. *huFNRG mice = humanized Fah^−/−^ NOD Rag1^−/−^ Il2rg^null^ mice, mpPHHs = mouse-passaged primary human hepatocytes*.

We quantified intrahepatic HBV DNA levels in isolated mouse-passaged primary human hepatocytes (mpPHHs) from the infected huFNRG mice and found 4.2-fold increased HBV-DNA levels in BCP-mutant infection compared to WT (normalized to total DNA) (**Fig. 4F**). Increased HBc expression in BCP-mutant infected mice was confirmed by western blot of isolated mpPHHs (**Fig. 4G**).

We also performed stable isotope labeling by amino acids via mass spectrometry (SILAC) in cultured mpPHHs to distinguish between increased viral antigen accumulation (“light amino acid-labeling”) and increased *de novo* production (“heavy amino acid-labeling”). This revealed a 4-fold increase in accumulated HBc protein in BCP-mutant infected hepatocytes compared to WT (**Fig. 4H**, p=0.0014) and 2.8-fold increase of HBs protein (**Fig. 4I**, p=0.0508). Quantification of newly synthesized proteins also trended towards higher *de novo* HBc and HBs synthesis in BCP-mutant infection than WT and PC-mutant infected mpPHH cultures (**Fig. 4K,L**). Precore protein abundance was comparable in WT and BCP-mutant infected cells and absent in PC-mutant infected hepatocytes (**Fig. 4J,M**).

Altogether, these data show that the BCP mutation in this HBV-D sequence increased viral replication and protein production compared with its parental WT sequence and isogenic PC-mutant.

### The BCP mutation remodels the infected hepatocyte proteome toward an inflammatory and carcinogenic signature

Given this striking increase in HBV-D BCP replication and viral protein expression, we also examined possible effects on the human hepatocyte proteome. In human hepatocyte chimeric mouse livers, the high sequence homology between human and murine proteins limits reliable species discrimination using peptide-based mass spectrometry. SILAC-based proteomics using cultured mpPHHs isolated from HBV-infected huFNRG mice circumvents these limitations because the cultures are highly enriched for human cells, the vast majority of which express HBV.[35,47,49]

Unbiased principal component analysis revealed distinct clustering of WT-, BCP- and PC-mutant-infected mpPHH cultures (**Fig. 5A**). Pairwise comparisons revealed many proteins altered between WT, PC-mutant and BCP-mutant-infected hepatocytes, with the largest differences between PC-mutant and BCP-mutant infected cells (**Fig S11**). A heatmap of all 906 differentially expressed proteins across all 3 conditions (WT, PC-mutant, BCP-mutant) showed all 6 possible clusters representing different expression patterns (**Fig. 5B**). Kyoto Encyclopedia of Genes and Genomes (KEGG) pathway analysis comparing which up- or downregulated proteins in BCP- and PC-mutant infected cells relative to WT revealed shared and distinct pathways (**Fig. 5C-F, Tables S5A-D**). Examples of shared KEGG pathways between BCP-mutant infection and PC-mutant infection were “Metabolic pathways” (upregulated in BCP- and PC-mutant vs. WT) and “Protein processing in endoplasmic reticulum” (downregulated in BCP- and PC-mutant vs. WT) (**Fig. 5C-F**).

**Figure 5:**
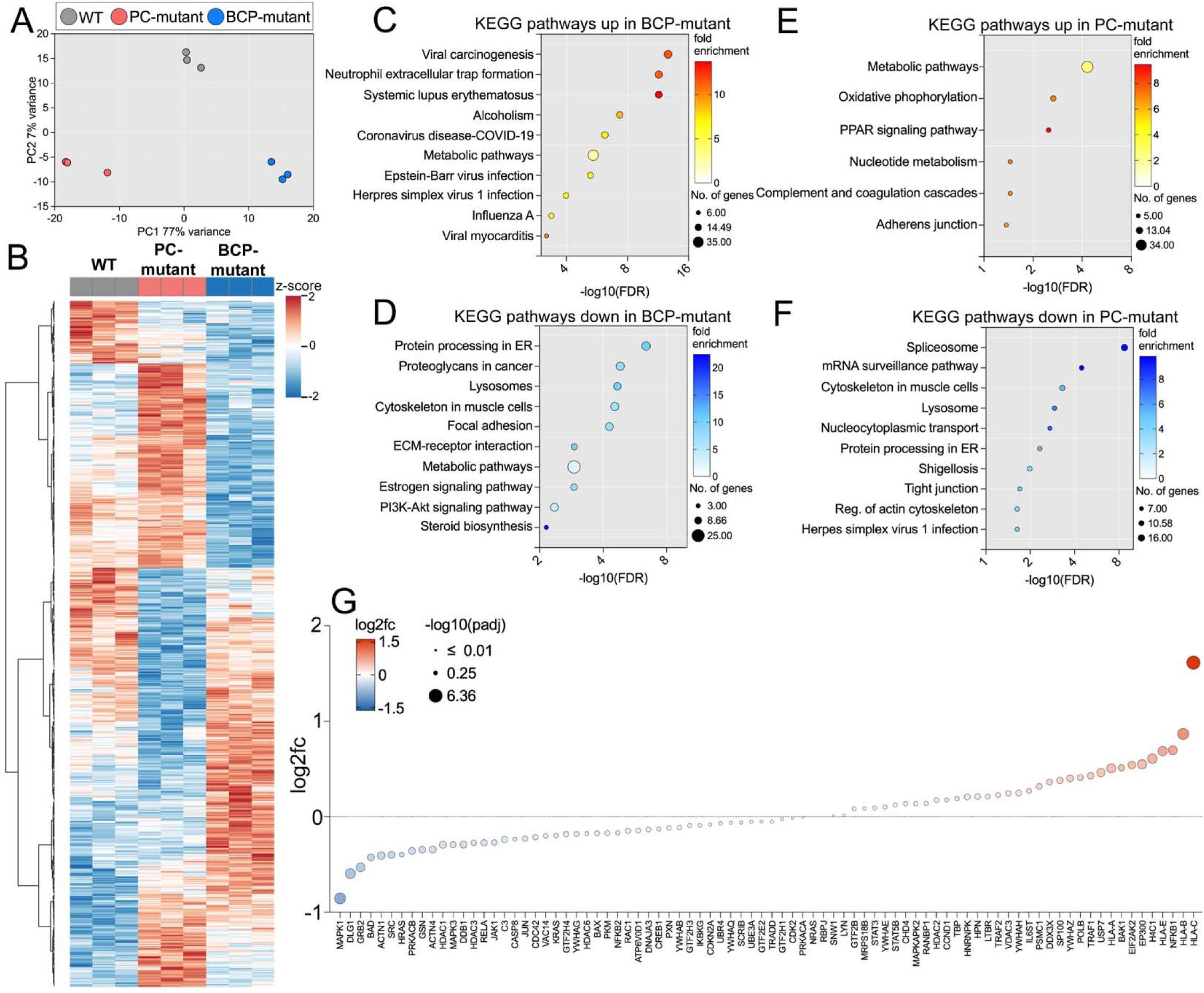
Distinct proteomic responses to HBV-D infection with WT, BCP-, or PC-mutants in mpPHHs. **(A)** Principal component analysis of protein expression in HBV-infected mpPHHs with HBV-D WT, BCP-mutant, or PC-mutant infection using all identified proteins. **(B)** Heatmap showing differential protein expression in cultured mpPHHs (all proteins with significant differential expression (p-adj < 0.05) in any pairwise comparison. **(C - F)** Enriched KEGG pathways in **(C)** upregulated proteins in BCP-mutant vs. WT-infection, **(D)** downregulated proteins in BCP-mutant vs. WT-infection, **(E)** upregulated proteins in PC-mutant vs. WT-infection, and **(F)** downregulated proteins in PC-mutant vs. WT-infection based on all significantly up- or downregulated proteins with a -log10 (p-adj) > 2. **(G)** Pairwise comparison of protein expression between BCP-mutant and WT-infected mpPHHs. Shown are all 84 proteins from the KEGG “Viral carcinogenesis” pathway detected via mass spectrometry. Positive log2fc-values indicate enrichment in BCP-mutant infection. *fc = fold change, FDR = false discovery rate, KEGG = Kyoto Encyclopedia of Genes and Genomes*.

The three pathways most significantly induced by the BCP-mutant compared to WT included “Systemic lupus erythematosus” and “Neutrophil extracellular trap formation”, which include many proteins involved in inflammation, and “Viral carcinogenesis” (**Fig. 5C**). Focusing on the 205 proteins in the “Viral carcinogenesis” KEGG pathway, 84 were differentially expressed in BCP-mutant mpPHHs compared to WT. These included upregulation of *HLA-A/B/C*, *HLA-E*, *SP100*, *EP300*, and *IL6ST*, as well as *TRAF1/2* and *NF-κB*, indicating an activation of inflammatory pathways in BCP-mutant infected hepatocytes. Furthermore, *USP7*[50], *EIF2AK2*[51], and *YWHAZ*[52] were also upregulated in BCP-mutant infection and have previously been associated with human HCC (**Fig. 5G**).

These data show that both the BCP and PC mutations perturb the proteome in human hepatocytes, with inflammatory and cancer-related pathway induction prominent in BCP-mutant infection.

### BCP-mutant infection drives a divergent, carcinogenesis-associated transcriptome that aligns with a poor-prognosis HBV-HCC cluster

The clinical association of BCP-mutants with HCC development[13–15], together with mutant-specific proteomic alterations in the “Viral carcinogenesis” pathway, prompted further investigation into the transcriptomic landscape in huFNRG mouse livers and public HCC datasets.

BCP-mutant mice clustered away from WT and PC-mutant animals on principal component analysis (**Fig. S12**) and induced the most transcriptional changes of the three HBV-D variants compared to uninfected livers (**Fig. 6A**). There was a strong correlation in gene expression between BCP-mutant vs. WT and BCP-mutant vs. PC-mutant infections (Spearman r = 0.69, p < 0.0001) (**Fig. 6B**) and transcriptional changes were consistent between individual infected mice (**Fig. 6C**). Collectively, these data indicate that the HBV-D BCP-mutant induces a transcriptomic profile that is divergent from WT and PC-mutant infection.

**Figure 6:**
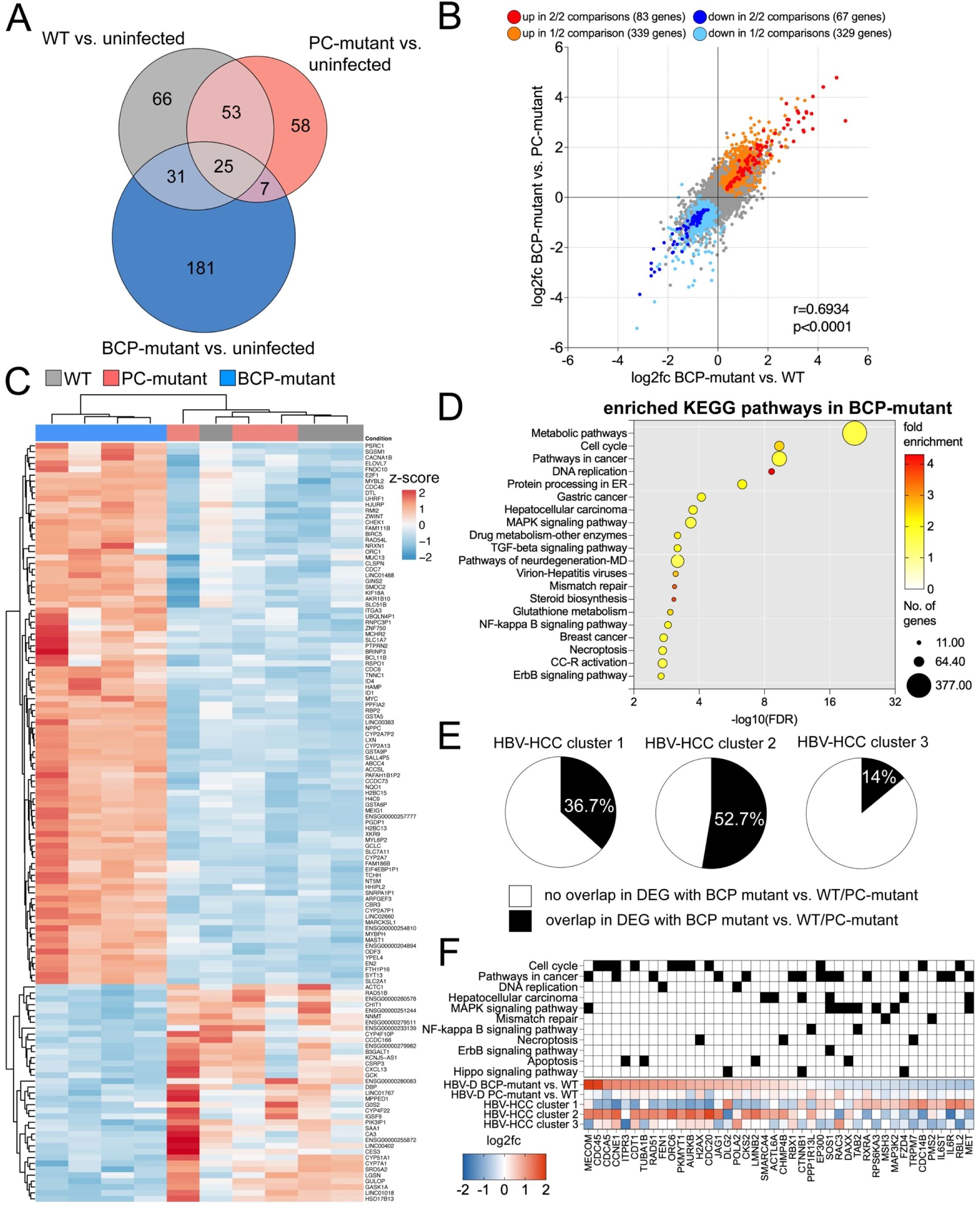
Transcriptomic analysis in HBV-D links the BCP mutation to a distinct cluster of HBV-associated HCCs. **(A)** Venn diagram of differentially expressed genes (DEGs, p-adj < 0.05) between uninfected huFNRG mice and mice infected with HBV-D WT, BCP-mutant, or PC-mutant virus. **(B)** Scatter depicting differential gene expression between BCP-mutant and WT infection infection (x-axis) and BCP-mutant and PC-mutant infection (y-axis) in huFNRG mouse livers. **(C)** Heatmap showing differential gene expression in livers from huFNRG mice infected with HBV-D WT, BCP-mutant, or PC-mutant virus (p-adj < 0.01, log2fc > |1.5|). **(D)** KEGG pathway enrichment analysis for BCP-mutant-associated DEGs compared to WT and PC-mutant infection. **(E)** Overlap of genes that show a significant differential gene expression (p-adj < 0.05) in HBV-D WT or PC-mutant vs. BCP-mutant infection in huFNRG mice and three distinct clusters of HBV-associated HCCs. **(F)** Gene expression comparison of HBV-D WT vs. BCP-mutant, HBV-D WT vs. PC-mutant, and HBV-HCC clusters 1-3. Filled black boxes in the upper panel indicate affiliation with KEGG pathways. All presented genes show significant (p-adj < 0.001) differential expression in two out of three HBV-HCC clusters. *CC-R = chemical carcinogenesis-receptor, DEGs = differentially expressed genes, ER = endoplasmic reticulum, fc = fold change, HCC = hepatocellular carcinoma, huFNRG mice = humanized Fah^−/−^ NOD Rag1^−/−^ Il2rg^null^ mice, KEGG = Kyoto Encyclopedia of Genes and Genomes*.

We next examined enriched KEGG pathways in BCP-mutant infection compared to isogenic WT or PC-mutant infection (**Fig. 6D, Table S6**). For BCP-mutant infection, most genes grouped in “Metabolic pathways”, with a total of 377 out of 1556 genes being differentially expressed. In addition, other metabolic and inflammatory pathways (e.g., “Virion-Hepatitis viruses”, “NF-kappa B signaling pathway”, “Necroptosis”) as well as multiple pathways linked to carcinogenesis (e.g., “Pathways in cancer”, “Gastric cancer”, “Breast Cancer”, “Chemical carcinogenesis-receptor activation”) and cell-fate regulation (e.g., “Cell-cycle”, “DNA replication”, “TGF-beta signaling pathway”) were enriched in BCP-mutant infection. Notably, the pathway “Hepatocellular carcinoma” showed significant enrichment, with 48 out of 148 constituent genes being differentially expressed. These analyses indicated that BCP-mutant infection caused transcriptomic changes associated with carcinogenesis.

We next examined whether the trajectories of dysregulated cancer-related genes in BCP-mutant infection converged those reported for human HBV-related HCCs (HBV-HCC). This comparison was complicated given the heterogeneous nature of HCCs.[53–55] Recently, Tian et al. described three distinct clusters of HBV-HCCs using transcriptomic data from The Cancer Genome Atlas (TCGA).[56,57] These HBV-HCC clusters were associated with differences in survival, with HBV-HCC cluster 2 exhibiting the poorest prognosis with activation of several cell cycle-related genes and cell death.

To assess a potential relationship between these three HBV-HCC clusters and BCP-mutant infection in huFNRG mice we then determined whether the 150 DEGs of the BCP-mutant compared to WT or PC-mutant infection were also differentially expressed across the three human HBV-HCC clusters. The greatest overlap was observed with HBV-HCC cluster 2 (52.7%), followed by cluster 1 (36.7%) and cluster 3 (14%) (**Fig. 6E**). Focusing on genes significantly dysregulated across the Tian HBV-HCC clusters, we found concordance in the direction of gene regulation upon BCP-mutant infection and HBV-HCC cluster 2. In contrast, gene regulation patterns appeared largely opposite in cluster 1, and no clear pattern was evident for HBV-HCC cluster 3 or changes induced by PC-mutant infection compared to WT (**Fig. 6F**).

These results show that for this HBV-D sequence, infection with BCP-mutant leads to the greatest disruption in the hepatocyte transcriptome compared to WT or PC-mutant infection. In BCP-mutant-infected hepatocytes, a substantial number of transcriptomic changes are enriched in carcinogenesis pathways, most consistent with HBV-HCC cluster 2 based on the Tian classification.[56]

### WT- and BCP-mutant HCCs cluster with tumor subtypes

The observed correlation between BCP-mutant induced transcriptomic changes and Tian HBV-HCC cluster 2 prompted us to investigate whether human HBV-HCCs arising in BCP-mutant infected individuals might exhibit distinct features compared to those arising in WT infection. We accessed transcriptomic data from all available HCC samples within TCGA dataset, using HBV genotypes A-E to screen for HBV transcripts.[41] HBV-RNA was found in 25.1% (93/371) of all HCC cases in TCGA (**Fig. S13A**), including 5 samples with no clinical HBV annotation.

We classified HBV-HCC cases as WT, BCP-mutant, PC-mutant, or double-mutant based on the predominant viral sequence at the BCP (nt1762/1764) and PC (nt1896) loci and considered cases as “mixed” samples if they contained ≥ 2.5% of both WT and mutant reads. In 31.3% of samples, the BCP-mutant was predominant (“BCP-mutant” or “BCP-mutant mixed”), and notably, almost half (45.3%) of the HBV-HCCs were generally classified as “mixed” (**Fig. 7A**). An illustration of relative abundance of WT or mutant reads at the BCP or PC loci per sample is shown in **Fig. 7B and Fig. S13B**, highlighting that heterogenous viral populations are present in a substantial fraction of HBV-HCCs.

**Figure 7:**
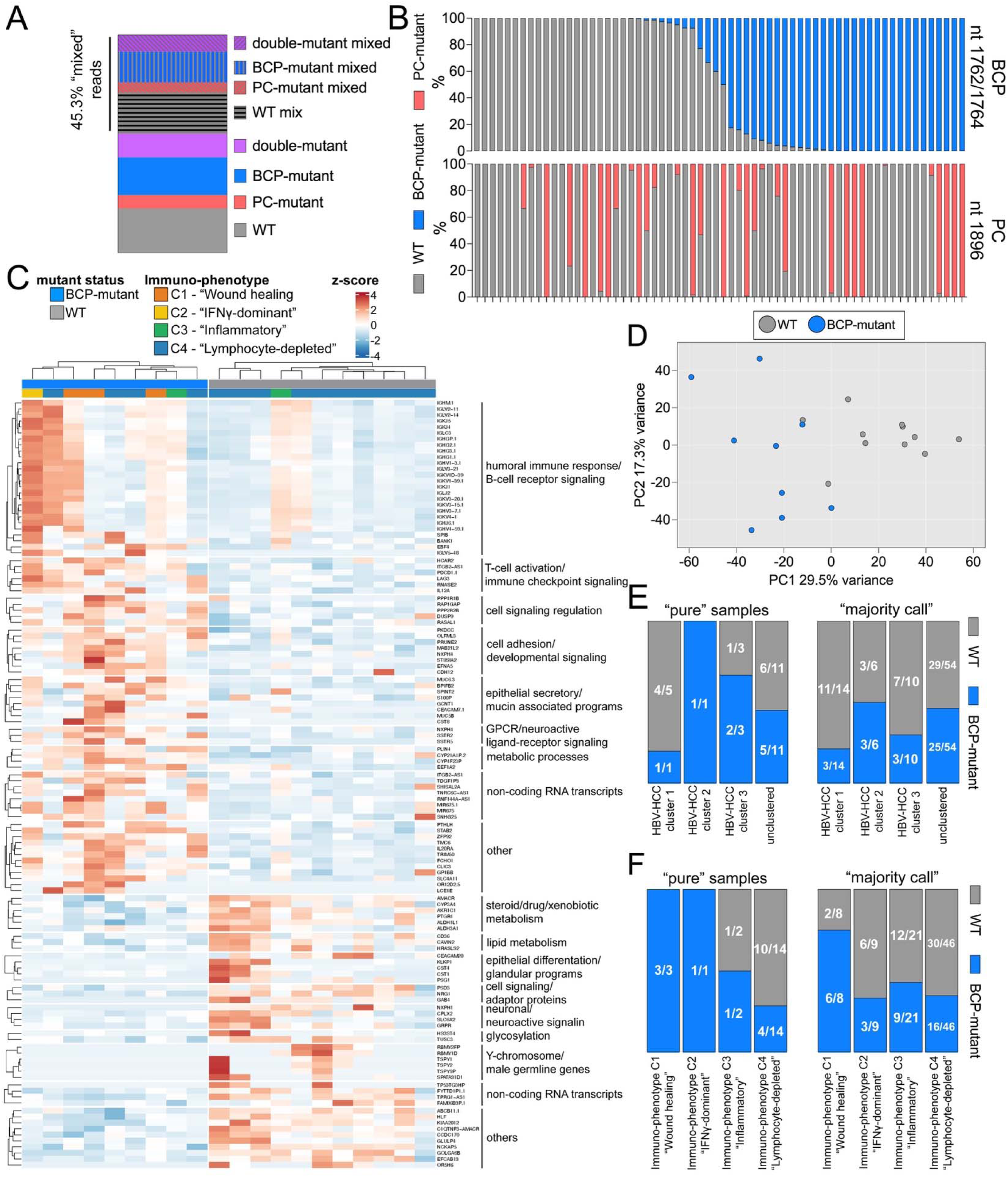
HBV mutant analysis in HBV-associated HCCs from the TCGA dataset reveals viral mosaicism and mutant-specific clustering. **(A)** Relative frequency of predominant viral sequences in TCGA HCC samples by HBV mutation. Samples were classified as “mixed*”* if the predominant sequence comprised < 97.5% of reads or if ≥ 2.5% of reads within the same sample harbored the alternate mutation (BCP or PC). **(B)** Relative abundance of WT or mutant sequences in HBV-RNAs detected in HBV-HCCs at BCP (nt1762/1764, upper panel) or PC (nt1896, lower panel). Only samples with sufficient coverage at both positions are shown here. Aligned columns represent one HCC sample. **(C)** Heatmap showing differentially expressed genes (p-adj < 0.0001, log2fc > |2|) between WT- and BCP-mutant-associated HCCs (“pure” HBV-HCC samples). **(D)** Principal component analysis of “pure” HBV-HCCs with WT or BCP-mutant infection based on the 500 most variable genes. **(E and F)** Association of HBV WT- and BCP mutant-associated HCCs with the HBV-HCC clusters from Tian et al. **(E)** or immune subtypes **(F)**. “pure” HBV-HCC samples: ≥ 97.5% WT or BCP-mutant reads at BCP loci 1762/1764 and 0 PC-mutant reads at PC locus nt1896 (samples with coverage < 10 at nt1896 excluded). “majority-call” HBV-HCC samples: ≥ 50% WT or BCP-mutant reads at BCP loci nt1762/1764 and ≤ 50% PC-mutant reads at PC locus nt1896. *fc = fold change, HCC = hepatocellular carcinoma, TCGA = The Cancer Genome Atlas*.

Using a strict filtering strategy that excluded all samples with a detectable PC mutation, we selected “pure” WT (n = 11) or BCP-mutant (n = 9) samples for comparison. A total of 1267 genes were differentially expressed, and the most significantly altered genes are shown in **Fig. 7C**. WT or BCP-mutant-associated HCC transcriptomes clustered distinctly (**Fig. 7D**). The BCP-mutant infected cohort exhibited an upregulation of immunoglobulin variable regions (numerous heavy- and light-chain segments) as well as B-cell markers (*SPIB, BANK1*) and markers of T-cell exhaustion (*PDCD1, LAG3*), suggesting increased immune cell infiltration. In addition, hepatocyte-specific metabolic markers (*CYP3A4, AKR1C1, ALDH3A1, ALDH1L1, CD36, CAVIN*) were downregulated while ductal and inflammatory epithelial markers (*MUC5B, MUC6, CEACAM7*) and stromal genes (*STAB2*) were upregulated, suggesting a relative loss of hepatocellular differentiation. We performed a similar analysis for all “pure” PC-mutant samples (n = 4) and found simultaneous upregulation of *LCN2, PLA2G2A, IFI44L*, and *XAF1*, consistent with an injury-reactive inflammatory tumor with activated interferon signaling (**Fig. S14**).

Hypothesizing that the BCP-mutation might affect the immunogenic phenotype of HCCs, we assessed the distribution across an established classification of immune subtypes of cancer.[58] Given the low number of samples with “pure” BCP-mutant reads, we also applied an approach that included samples segregated by the predominant read at the BCP locus (“majority call”) (**Fig. S15**). To complement the overlap in DEGs between huFNRG mice and reported human HBV-HCC clusters, we first examined the clustering of BCP-associated HBV-HCCs as reported by Tian et al. and, again, found a trend toward enrichment for cluster 2 (**Fig. 7E**). Among HBV-HCCs classified as “pure”, 3/3 HBV-HCCs of the C1 immune-phenotype (“Wound healing”) subtype were BCP-mutant-associated, and 10/11 WT-associated HCCs were assigned to subtype C4 (“Lymphocyte depleted”). In HBV-HCCs classified by “majority call”, BCP-associated HCCs were again enriched in subtype C1 (6/8), whereas WT-associated HCCs were more frequently assigned to subtypes C2 (“Interferon-gamma dominant”), C3 (”Inflammatory”), and C4. No HBV-HCCs with detectable HBV RNA were clustered into subtype C5 (“Immunologically quiet”) or C6 (“TGF-beta dominant”) (**Fig. 7F**).

Taken together, these analyses demonstrate that HBV-HCCs harbor considerable viral heterogeneity, but they cluster by the dominant HBV species in the sample (WT, PC- or BCP-mutant) and WT- and BCP-mutant-associated HCCs exhibit distinct tumor-associated transcriptomic and immunological profiles.

## Discussion

Many studies have linked the BCP mutation to worsened clinical outcomes, including increased HCC risk.[9–15] Since HBV has long been considered non-cytopathic, hepatic injury occurring in chronic HBV infection is attributed to host immune-mediated mechanisms.[16–19] However, studies in immunodeficient hepatocyte chimeric mice challenge this paradigm, suggesting that HBV may acquire cytopathic features in the context of BCP and PC mutations.[27–29] Most recently, Uchida et al. observed accelerated viremia and loss of engrafted human hepatocytes upon infection with a PC/BCP-double-mutant HBV-C isolate.[29] These studies were constrained by reliance on patient-derived viral stocks, which precluded linking phenotypes to specific viral mutations though Uchida et al. addressed this by generating an isogenic control in cell culture.[29]

To overcome these challenges, we generated infectious viral stocks in huFNRG mice via intrahepatic injection of rcccDNA. This approach reliably produced infectious stocks of WT and isogenic BCP- and PC-mutants without evidence of spontaneous acquisition of these mutations or reversion to WT in any of the sequences tested (**Fig. 1, Fig. S1** and **Fig. S2**). BCP-mutants in fact exhibited higher replication than WT in 2/3 sequences tested, demonstrating enhanced replicative fitness in the absence of immune pressure (**Fig. 2**). These data support a model in which immune control, rather than intrinsic viral fitness, drives differences in viremia levels between HBeAg-positive and HBeAg-negative phases.[3]

The mechanisms by which the BCP-mutant shapes the hepatic environment remain incompletely understood. Given the accelerated viremia observed in BCP-mutant infection, it appears plausible that enhanced viral replication, together with viral protein accumulation contribute to intrinsic hepatocyte reprogramming. This notion is supported by reports of fibrosing cholestatic hepatitis in immunosuppressed liver transplant recipients with high levels of viral replication and viral protein accumulation.[26] Uchida et al. reported increased ER stress in a double-mutant HBV[29], which we did not observe. Nevertheless, they showed a similar fitness hierarchy (double mutant > PC-mutant > BCP-mutant > WT) to our results using another HBV-C sequence (**Fig. 2** and **Fig. S5**).

We observed that transcriptomic changes induced by BCP-mutant infection in huFNRG livers aligned with a distinct cluster of HBV-HCCs (**Fig. 6E,F**).[56] We interpret these overlaps as evidence of a hepatocyte-intrinsic signature that may contribute to specific HBV-HCC subtypes. To explore a link in human disease, we analyzed RNA-seq data from HBV-HCCs in TCGA, stratifying tumors by expression of WT or BCP-mutant HBV transcripts (**Fig. 7** and **Fig. S15**). This analysis suggested an increased immune-infiltration in BCP-mutant carriers. The observed divergence in immune phenotypes might prove relevant in the future as immunotherapeutic strategies for HCC evolve.[59] We furthermore identified heterogeneous viral populations within HBV-associated HCCs, raising intriguing questions about the coexistence, competition, and coevolution of HBV variants.

While cccDNA can be detected in HCCs[60], studies in hepatoma cells in vitro[61] and chimeric mice[62] have demonstrated that cccDNA is lost in proliferating cells. This raises a conceptual question regarding how BCP-mutation-driven hepatocyte reprogramming would be sustained in cells undergoing malignant transformation. Beyond cccDNA conservation in HCCs, several non-mutually exclusive mechanisms may contribute to the persistence of BCP-mutant-induced changes. One possibility is that BCP-mutant-driven alterations operate through a “hit-and-run” mechanism, whereby the mutant virus establishes a pro-oncogenic state that persists independently of continued viral replication. Additionally, in untreated patients, the cccDNA pool may be replenished through reinfection. Finally, HBV RNAs detected in HBV-HCCs may originate from integrated HBV DNA, thereby sustaining BCP-mutant transcript expression independently of cccDNA. Additional mechanisms contributing to the increased HCC risk in BCP-mutant carriers merit consideration. HBV-HCCs frequently harbor HBV-integration events, with approximately 40% of HBV breakpoints located in HBx near the BCP region[63] and the BCP mutation might also modulate the transcription of distinct HBx transcript variants.[64] Future studies using isogenic HBV strains in hepatocyte chimeric mice may help clarify how these candidate mechanisms differentially contribute to malignant transformation.

Several limitations of this study should be acknowledged. As we only studied one parental sequence per genotype, we cannot attribute any observations to inter-genotypic differences. Future studies across larger collections of HBV sequences and isolates may determine whether our findings can be replicated in other HBV-D sequences and other HBV genotypes. Differences between the transcriptomic data from bulk liver samples and proteomic data from mpPHHs might be driven by selection biases through the human hepatocyte isolation protocol and maintaining these cells in culture. In addition, differences between huFNRG cohorts used in separate experiments might contribute to transcriptomic differences observed between mice infected with different HBV WT sequences (**Fig. 3B**). Furthermore, the observed transcriptomic overlap between BCP-mutant-infected hepatocytes and HCC-associated gene signatures is correlative in nature and does not establish a causal relationship. Definitive causal determination would require experimental systems that currently do not exist, as PHH cannot be experimentally transformed to human HCCs, and available non-human HBV models carry translational limitations with regard to human BCP-driven carcinogenesis.[31,65,66]

In summary, this study provides a controlled isogenic framework that highlights divergent biology of the BCP- and PC-mutants in human hepatocyte chimeric mice. Our data indicate that the BCP mutation perturbs hepatocyte homeostasis via adaptive immune system-independent mechanisms, supporting a model of HBV-mutant-driven hepatocarcinogenesis. We also identify heterogeneous viral populations in HBV-associated HCCs and a BCP-mutant-associated HCC phenotype. Incorporating viral-variant biology into current immune-centric frameworks for chronic HBV is therefore essential to explain heterogeneous disease trajectories and to improve risk stratification, therapy, and surveillance strategies.

## Materials and Methods

### Animal studies

*Fah^-/-^*NOD *Rag1^-/-^ Il2rg^null^* (FNRG) mice were created by backcrossing *Fah^-/-^*liver injury mice[67] (provided by Markus Grompe, Oregon Health and Science University) to NOD *Rag1^-/-^ Il2rg^null^* (NRG, Jackson Laboratories) as previously described.[68] To create humanized (hu)FNRG mice, animals were preconditioned with retrorsine (catalog no. R0382; Sigma Aldrich) and withdrawn from nitisinone (Yecuris) prior to transplantation with 5×10^5^ primary human hepatocytes from donor HUM4188 or HUM4143 (Lonza).[35] Animals were cycled on nitisinone for 10-12 weeks. Human engraftment was confirmed using a species-specific Enzyme-linked immunosorbent assays (ELISA) to quantify serum human albumin levels (Bethyl Labs). For infection experiments, only highly humanized mice with serum human albumin levels of approximately 10mg/ml were used. For intrahepatic injection of recombinant covalently closed circular HBV DNA (IHI rcccDNA), animals with serum human albumin levels >0.5mg/ml were used. Infection experiments were performed in highly humanized huFNRGs with serum human albumin levels around 10mg/ml and randomization via drawing lots was used for allocation of animals.[35] Histological analysis of tissue was performed blinded. To avoid cross contaminations, no blinding was exercised during the conduct of the experiment. Isolation of mouse-passaged primary human hepatocytes (mpPHHs) from huFNRG mice was performed as previously described.[35] All mice used in this study were housed and bred at The Rockefeller University Comparative Biosciences Center in accordance with the NIH Guide for the Care and Use of Laboratory Animals. All procedures were approved by the Rockefeller University IACUC (protocol #18063, #21056 and #24022). Statistical analyses and data visualization were performed using GraphPad Prism version 11.0.2. (GraphPad Software, San Diego, CA, USA).

### Viral stock preparation and intrahepatic injection of recombinant ccc-like HBV DNA

RcccDNA DNA for intrahepatic injection was generated based on a previously described method by Lempp et al.[38] In brief, full-length 3.2kb HBV DNA sequences were enzymatically digested from a specifically designed plasmid DNA construct and purified after gel electrophoresis (1% agarose gel) using the QIAquick® Gel Extraction Kit (Qiagen, catalog no. 28704). NEB T4 Ligase (M0202M) and T4 DNA Ligase Reaction Buffer (B0202S) were added to achieve self-ligation after diluting the reaction with nuclease free water to a DNA concentration of 1-3ng/µl to prevent ligation between DNA molecules. Ligation was performed over night at room temperature and DNA was purified using the QIAquick® PCR Purification Kit (Qiagen, catalog no. 28106). DNA was then again run on a 1% agarose gel to purify self-ligated rcccDNA. Up to 2µg of HBV rcccDNA were used for IHI, which was performed under strictly sterile condition and isoflurane anesthesia. Following a right subcostal incision (∼1cm), the peritoneum was mobilized and opened. A total volume of 200µl of rcccDNA (diluted in phosphate buffered saline without calcium or magnesium, PBS^-/-^) was injected into the large liver lobe using a 34-gauge syringe (Hamilton company 7633-01& 207434-10) and 10-20 injections with 10-20µl each. After the injections, mild pressure was applied to the injection sites using sterile cotton swaps to achieve hemostasis. Afterwards, the peritoneum was closed with Vicryl sutures and the skin was adapted with metal clips. For local analgesia, bupivacaine was applied, and for general analgesia, mice received two doses of buprenorphine (0.05mg/kg body weight: Abbot Animal Health). Metal clips were removed 14 days after surgery.

### Hepatitis B virus sequencing and SNV calling

5µl of DNA extracted from serum was used for HBV DNA amplification with genotype specific primers producing a near full-length amplicon as previously described.[69] The amplification primers are listed in **Table S2**. PCRs were performed using the Roche Hifi Kit with the following amplification settings (initial denaturation for 3 minutes at 95°C, followed by 25 – 35 cycles of 95°C for 20s, 55°C for 15s and 72°C for 144s. Final extension at 72°C for 5 minutes). PCR products were purified from a 1% agarose gel with a column-based cleanup (Zymoclean Gel DNA Recovery Kit, catalog no. 11-301C). Amplicons were quantified by fluorometric measurement and submitted to the MGH DNA Core for library preparation and sequencing. Sequencing data was trimmed using the BBDuk Adapter/Quality Trimming tool (Version 38.84) in Geneious Prime® (2025.2.2) [settings: Kmer Length 27, minimum quality 25, minimum length 25, minimum entropy 0.1, window size 50, kmer size 5]. Reads were then mapped to the respective wild-type consensus sequence using the “find variations/SNP” workflow. SNVs were only counted in case of an average quality >25 and the absence of significant strand bias (strand bias >65% with p-value < 0.05). As positions proximal to the amplification primers are covered predominantly by reads originating in direction the respective primer, strand-bias was ignored within 60bp of the amplification primers. To define the minimum coverage required for reliable variant detection at each position, we applied an empirical model that accounts for theoretical sampling requirements and sequencing errors of Illumina MiSeq, taking previous reports and recommendations into account.[70–74] The required per-site depth for detecting a variant at frequency was defined by the formula below including a 2.5-fold safety margin to account for amplification and sequencing errors. SNVs with coverage below D_required_(p)were excluded.

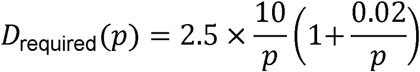

### Quantitative PCR analysis of HBV DNA

To quantify HBV DNA in the serum of huFNRG mice, 25µl of serum were diluted in 175µl of DPBS (Dulbecco’s phosphate buffered saline without calcium or magnesium, DPBS^-/-^) and DNA was extracted using the QIAamp DNA Blood Mini Kit (Qiagen, catalog no. 51106). DNA was eluted in 60µl elution buffer and HBV DNA levels were then measured by qPCR following the previously described method using a Taqman-based assay.[75] The qPCR primers for different HBV strains are listed in supplementary **Table S1**.

### HBeAg and HBsAg chemiluminescence immunoassay

Chemiluminescence Immunoassays (CLIA) for the detection of HBsAg and HBeAg in the serum of huFNRG mice were performed as previously described[76] per manufacturer’s instructions (DiaSino Co., China).

### Histology

Histological procedures were performed according to best practices.[77] Liver tissues were dissected to appropriate size and fixed in 10% neutral buffered formalin for ≥48 hours (Fischer Scientific, Cat # SF100-4, Lot# 216889). Tissue was processed in Sakura VIP6 processor following a one hour, 13 step program and paraffin-embedded immediately upon completion of processing. Five-micron tissue sections were collected onto Plus slides (Epredia, Cat # TT-4041L-PS-8218-1), air-dried and stored at room temperature prior to use. Each sample was histochemically stained with Hematoxylin-Eosin for morphological evaluation[78]. Picrosirius red stain for collagen was performed according to the method of Puchtler[79]. In brief, sections were deparaffinized in xylene (3 changes), rehydrated through graded alcohols (3 changes 100% ethanol, 3 changes 95% ethanol) and rinsed in distilled water. Sections were stained for 1 hour in 1.0% Picric Acid Sirius Red (Abcam, catalog # ab246832, Lot # 1057287-2) solution at room temperature. Sections were washed in two changes of 0.5% glacial acetic acid in water, dehydrated with 2 changes of 100% ethanol followed by three changes of xylene and mounted with permanent media. Immunohistochemistry was performed on Ventana Medical Systems Discovery Ultra platform using Ventana reagents unless otherwise specified. Mouse anti-human unconjugated Hepatitis B Virus Surface Antigen (HBVs), clone S1-210 (Ventana Medical Systems Cat # 760-2647, Lot #31099 RRID, AB_2335993), polyclonal rabbit anti-human unconjugated Hepatitis B Virus Core Antigen (HBcAg) (Ventana Medical Systems Cat # 760-2646, Lot #17099, RRID, AB_2335959) and polyclonal rabbit anti-human unconjugated Fumarylacetoacetate Hydrolase (FAH) (Yecuris Cat # 20-0034, Lot# LXA00003, RRID unassigned) protocols were optimized on normal and known HBV core and surface positive samples. In brief, sections were deparaffinized online at 69°C for 24 minutes (Ventana Discovery Wash Cat# 950-510). HBVs was antigen retrieved at 91°C for 32 minutes (Ventana Ultra CC2 Cat# 950-223) and HBcAg was retrieved at 95°C for 56 minutes (Ventana Ultra CC1 0Cat# 950-224). FAH did not require antigen retrieval. Endogenous peroxidase was blocked on all samples for 4 minutes. Anti-HBVs was applied neat and incubated for 60 minutes at 37°C. Anti-HBcAg was applied neat and incubated for 8 hours at room temperature. Anti-FAH was diluted 1:800 (Ventana Cat # ABD250) and incubated for 3 hours at room temperature. Primary antibody was detected using goat anti-rabbit (Ventana Cat # 760-4311) or goat anti-mouse (Ventana Cat # 760-4310) horseradish peroxidase conjugated multimer incubated for 8 minutes, respectively. The complex was visualized with 3,3 diaminobenzidene and enhanced with copper sulfate. Slides were washed in distilled water, counterstained with hematoxylin, dehydrated and mounted with permanent media. Negative controls consisted of diluent only. All slides were scanned on an Aperio (Leica) AT2 captured at 40x magnification and visualized on eSlideManager software (Version 12.5.0.6145). Image analysis was performed in Aperio ImageScope (v12.4.6.5003) using the Positive Pixel Count v9 algorithm and QuPath[80] (v0.6.0).

### Western Blotting

PHHs were isolated from infected huFNRG mice as described above and 10^6^ isolated mpPHHs were used for protein extraction. Cells were lysed in an IGEPAL-lysis buffer (1% IGEPAL-CA630 (NP-40, Sigma, 18896), 50mM Tris-HCl at pH 8.0, 1mM EDTA, 1x Proteinase inhibitor, Roche 11697498001). The lysate was then mixed with 6xSDS sample buffer (300mM Tris (pH 6.8), 60% glycerol, 12% SDS, 0.2% bromophenol blue, 20% (v/v) β-mercaptoethanol), boiled for 10 mins, placed on ice for 15 mins, and then separated by SDS-PAGE on a precast 4-20% gradient polyacrylamide gel (4-20% Mini-PROTEAN TGX Precast Protein Gels, 10-well, 30ul, Bio-Rad #4561093) in 1xSDS-PAGE running buffer (25mM Tris base, 250mM glycine, 0.1% SDS). Gels were transferred onto a Polyvinylidene difluoride (PVDF). The gel was soaked in transfer buffer (25mM Tris base, 250mM glycine, 20% methanol) for 20 mins. The PVDF membrane was washed in 100% methanol for 15s, then in ddH2O for 2 mins before moving to transfer buffer for 15 mins. Transfer was run for 2h at 400mA (Mini Trans-Blot Electrophoretic Transfer Cell, #1703930) in transfer buffer. The membrane was then immersed in 100% methanol for 15s, rinsed twice with ddH2O, and washed in PBST (Phosphate-Buffered Saline with Tween-20) before adding blocking solution (PBST with 5% nonfat milk) for 1h at room temperature. The Blocked membrane was incubated with the respective antibodies (HBcAg: mouse MAb T2221 2AHC24, Tokyo Future Style, Japan) in incubation buffer (PBST with 1% nonfat milk) overnight at 4 °C. Horseradish peroxidase-conjugated Rabbit anti-mouse (Pierce 31452) was used as the secondary antibody and added after three washes with PBST and incubation at room temperature for 1h. Enhanced lumino-reagent and oxidizing reagent were combined at equal volume to make the chemiluminescent reagent mix. The membrane was placed in the imager, and the chemiluminescent reagent mix was pipetted over the membrane 1 min before imaging. After imaging, the exposure reagents were washed with PBST for 10 minutes at room temperature, 3 times. The membrane was then incubated with Secondary Antibody Anti-β-Actin-Peroxidase antibody (Mouse monoclonal, Millipore SiGMa A3854) solution and imaged again for detection of beta actin.

### Cell culture for pulse SILAC proteomics and sample preparation

Mouse-passaged primary human hepatocytes (mpPHH) were isolated from HBV-infected huFNRG mice as previously described in detail[35], plated in collagen-coated 12-well plates, and cultured for ≥7 days in a 5% CO_₂_ incubator. Cells were maintained in hepatocyte culture medium (HCM), which was refreshed every 2-3 days to ensure optimal viability. For stable isotope labeling, cells were first cultured for 24 h in SILAC RPMI 1640 medium (Thermo Fisher Scientific, Cat. # A33973) supplemented with L-lysine·2HCl and L-arginine·HCl (light SILAC medium). Cells were then cultured for an additional 48 h in SILAC RPMI 1640 medium supplemented with ^13^C_6_^15^N_2_ L-Lysine·2HCl and ^13^C_6_^15^N_4_ L-Arginine·HCl (heavy SILAC medium). At the end of the labeling period, cells were washed with ice-cold PBS, harvested using cell lifters, pelleted by centrifugation, and stored at −80°C until protein extraction. For LC/MS/MS analysis cell pellets were lysed in 8 M urea dissolved in 50 mM ammonium bicarbonate (ABC) buffer (pH 7.8) containing 1 % protease inhibitor cocktail (Sigma, P8340). Following sonication, protein concentrations were determined using a BCA assay kit (Thermo Fisher Scientific, 23225). Proteins were reduced with 10 mM dithiothreitol, alkylated with 50 mM iodoacetamide, and digested with Lys-C (Fujifilm Wako, 129-02541) for 1 hour at room temperature. Samples were then diluted 5-fold with ABC buffer and further digested overnight with trypsin (Promega, V5113). After digestion, samples were acidified with 10% trifluoroacetic acid to a final concentration of 0.5%. The resulting peptides were desalted using StageTips[81] packed with Empore SDB-XC (CDS, 98-0604-0224-9EA) as previously described.[82]

### LC/MS/MS analysis

LC/MS/MS analysis was performed using an Ultimate 3000 RSLC pump (Thermo Fisher Scientific) coupled to a PAL-xt HTC autosampler (CTC Analytics) and a timsTOF HT mass spectrometer (Bruker). Peptides were separated on an in-house packed needle column (100 μm i.d., 6 μm needle opening, 25 cm, Reprosil-Pur 120 C18-AQ 1.9 μm reversed-phase material; Dr. Maisch GmbH, r119.aq.) maintained at 50 °C. A 90-minute linear gradient from 4% to 32% acetonitrile containing 0.1% formic acid was applied at a flow rate of 500 nL/min. The trapped ion mobility spectrometry (TIMS) section was operated with a ramp time of 100 ms and a scan range of 0.6–1.5 Vs cm^-2^.[83] MS/MS spectra were acquired in diaPASEF mode, 1 MS1 scan followed by 25 dia-PASEF MS/MS scans with an estimated cycle time of 2.76 seconds, using isolation windows as previously described (**Table S7**).[84] The collision energy was set to 20 eV at 0.6 Vs cm^-2,^ increased linearly to 59 eV at 1.5 Vs cm^-2^.

### Proteomics Database search and Data analysis

Raw data were processed using Spectronaut (ver. 20.1) in directDIA (library-free) mode. The search was performed against the Swiss-Prot human database (ver. 2025_03) supplemented with genotype DE19 hepatitis B virus sequences using strict Trypsin/P specificity with allowance of up to two missed cleavages. Carbamidomethylation of cysteine was set as a fixed modification, while acetylation of protein N-termini and oxidation of methionine were set as variable modifications. Lys8 and Arg10 were specified for Channel 2 in the “Labeling” section. PSM, peptide, and protein identifications were filtered at false discovery rate of 1 % in the Pulser search. The library containing both light and heavy labeled peptides was constructed with the “In-Silico Generate Missing Channels” function. Quantification was performed for the identified precursors using “Group Qvalue” function within the “Multi Channel Qvalue Filter”. The resulting protein table was used for quantitative analysis. MS2 intensities from the heavy channel (PG.MS2Channel2) were used to quantify newly synthesized proteins. Proteins with missing values in any of the 18 samples were excluded from further analysis. Protein abundance values were normalized using the variance stabilizing normalization (VSN) method implemented in the vsn R package (ver. 3.74.0). Principal component analysis (PCA) was performed using the scikit-learn package (ver. 1.5.1) in Python. Differentially expressed proteins were identified using the moderated t test of Limma package in R (ver. 3.62.2)[85]; contrasts are detailed in **Table S8**. Proteins with Benjamini-Hochberg adjusted *p*-values (q-values) < 0.01 were considered significantly regulated in each contrast. KEGG pathway enrichment analysis was performed using ShinyGO v0.85 in reference to the human genome (GRCh38.p14). Heatmaps were generated using the seaborn Python package (ver. 0.13.2). Hierarchical clustering was performed using cosine distance and Ward’s linkage method with the Scipy library (ver. 1.14.0). The mass spectrometry raw data have been deposited in the jPOST repository[86,87] with the dataset identifier PXD072535 (JPST004281).

### RNA-sequencing and bioinformatics analysis

Bulk liver tissue (<25mg) or isolated human hepatocytes (10^6^ cells) were harvested at the individual peak viremia level of HBV infected huFNRG mice or respective time points in uninfected mice. The samples were homogenized using glass-beads (1mm, Biospec, catalog no. 11079110) in TRIzol® Reagent (Invitrogen, catalog no. 155966018). Total RNA was extracted via Chloroform extraction using MaXtract^TM^ High Density Columns (Qiagen, catalog no. 129056) and RNeasy® Mini Kit (Qiagen, catalog no. 74104) including on-column DNase treatment (Qiagen, catalog no. 79254) according to manufacturer’s instructions. RNA was eluted in 50µl RNase-free water and RNA concentration, and purity were assessed by Nanodrop (A260:A280 ratio 1.8 – 2.2). Samples were submitted to GENEWIZ (Azenta Life Sciences) for library preparation and sequencing (minimum RNA integration number (RIN) > 6.0). mRNA libraries were prepared using poly(A) enrichment following the service provider’s standard protocol. Sequencing was performed using Illumina NovaSeq 2×150 bp, with a targeted depth of 50 million reads per sample. RNA-seq reads were aligned using STAR (v2.4.2a) separately for human and mouse transcripts. Reads were first aligned to the human reference genome (GRCh38) using STAR mismatch and minimum alignment length filters (*--outfilterMismatchNoverLmax 0.04*, *--outFilterMatchNmin 20*), and read counts were generated using STAR *--quantMode GeneCounts*. Unaligned reads were exported using *--outReadsUnmapped Fastx* and subsequently aligned to the mouse reference genome (mm10) with unique-mapping enforced with --outFilterMultimapNmax 1. Gene counts generated by STAR were aggregated and used for differential expression analysis using DESeq2. Prior to differential expression analysis, genes with very low expression in more than 80% of samples were filtered out. Genes were considered differentially expressed if they met both statistical significance (p <= 0.05 and |log 2 fold change| >= 1.5) thresholds, unless stated otherwise.

### TCGA analyses

Aligned .bam files for each HCC sample were accessed via the TCGA Data Commons portal. Per position viral genotypes were called via bam-readcount[88] with default settings for the HBV reference strain AF121243.1. Mean coverage was estimated as the total nucleotide coverage from MAPQ ≥30 reads divided by the length of the viral contig. Processed RNA-seq counts were fetched from the recount3[89] server and analyzed for principal component analysis and differential gene expression as described for the murine data. For BCP mutant analysis, differentially expressed genes were computed using DESeq2[90] and principal components were computed based the top 500 most variables genes among top 2000 differentially expressed genes between BCP-Mutant and wild-type samples. Clustering statistical significance is then determined based on Mann-Whitney U test of the second principal component. For subtype enrichment analysis, fisher exact test was performed based on HBV-HCC cluster assignments obtained from Tian et al and the immune cluster assignment retrieved from Thorsson et al. The differentially expressed genes heatmap was generated via the ComplexHeatmap package[91] and survival analysis were performed via the “survminer” R package.[92]

## Author Contributions

Conceptualization, L.L.S., C.M.R., E.M., Y.P.J.; investigation, L.L.S., G.D., H.C., Y.Y., K.O., X.H., M.Z., Y.Z., C.Z., A.R., Ev. M., C.A.F., A. A., L.G., L.C., C.F., H.H., M.K., C.L., E.M., Y.P.J; supervision, C.M.R., A.S.L., W.M.S., E.M., Y.P.J; visualization, L.L.S., G.D., H.C., M.Z., K.O., H.H.; E.M., Y.P.J; writing—original draft, L.L.S., E.M., Y.P.J,; writing—review & editing, L.L.S., G.D., H.C., Y.Y., K.O., X.H., M.Z.,, Y.Z., C.Z., A.R., C.Q., E.Mo., C.A.F., A.A., L.G., L.C., C.F., H.H., M.K., A.S.L., C.L., W.M.S., C.M.R., E.M., Y.P.J.

## Data availability statement

Data are available upon reasonable request. Materials generated can be shared through material transfer agreements in compliance with our institution’s policies.

## Research Ethics Approval: Human Participants

This study does not involve human participants. Human transcriptomic data were obtained from The Cancer Genome Atlas (TCGA) Research Network and analyzed as publicly available, de-identified data.

## Research Ethics Approval: Animals

This study involves animals (mice) and was approved by The Rockefeller University IACUC (protocol #18063, #21056 and #24022).

## Supplementerary Materials

**Supplementary Fig. 1:** Successful launch of HBV strains and isogenic variants via IHI rcccDNA across genotypes A–D and patient consensus sequences.

**Supplementary Fig. 2:** Viral sequencing of IHI rcccDNA launches and infection experiments.

**Supplementary Fig. 3:** Hepatitis B infection with isogenic variants in HBV-A, HBV-C, and HBV-D: single mouse viremia.

**Supplementary Fig. 4:** BCP-mutant infection consistently accelerates viremia kinetics in HBV-D infection.

**Supplementary Fig. 5:** The double-mutant variant carrying both the PC-and BCP-mutation accelerates viral kinetics in HBV-C.

**Supplementary Fig. 6:** Time course of serum HBeAg and HBsAg levels in HBV-A, HBV-C, and HBV-D infection.

**Supplementary Fig. 7:** The human graft remains stable huFNRG mice with HBV infection.

**Supplementary Fig. 8:** Liver histology in HBV WT infection.

**Supplementary Fig. 9:** Proportion of human-, murine-, and ambiguously mapped reads in huFNRG-derived RNA.

**Supplementary Fig. 10:** Picrosirius red-staining in HBV-D infection.

**Supplementary Fig. 11:** Volcano plots of differentially expressed proteins identified by SILAC-based proteomics in pre-infected mpPHHs.

**Supplementary Fig. 12:** Principal component analysis of RNA-seq data from bulk liver tissue of huFNRG mice infected with HBV-D WT, BCP-mutant, or PC-mutant.

**Supplementary Fig. 13:** HBV sequencing coverage and relative variant abundance in human HCC.

**Supplementary Fig. 14:** Comparison of HBV-associated HCCs from the TCGA dataset: PC-mutant vs. WT.

**Supplementary Fig. 15:** Comparison of HBV-associated HCCs from the TCGA dataset: BCP-mutant vs. WT (“majority call”).

**Supplementary table S1:** HBV qPCR primer list

**Supplementary table S2:** HBV sequencing primer list

**Supplementary table S3:** Serum HBeAg levels in huFNRG mice

**Supplementary table S4:** Serum HBsAg levels in huFNRG mice

**Supplementary table S5:** Dysregulated KEGG pathways in SILAC proteomics in HBV-D infected mpPHHs

**Supplementary table S6:** KEGG pathways enriched in HBV-D BCP-mutant infected huFNRG mice (RNAseq)

**Supplementary table S7:** SILAC proteomics - PASEF acquisition table

**Supplementary table S8:** SILAC proteomics - contrasts

## Abbreviations

BCP: basal-core promoter
CLIA: Chemiluminescence Immunoassays
DEG: differentially expressed gene
DM: double mutant
DNA: deoxyribonucleic acid
ELISA: enzyme-linked immunosorbent assay
HBeAg: hepatitis B e antigen
HBsAg: hepatitis B s antigen
HBV: hepatitis B virus
HCC: hepatocellular carcinoma
H&E: hematoxylin and eosin
huFNRG mouse: humanized *Fah^−/−^* NOD *Rag1^−/−^ Il2rg^null^* mouse
IHI: intrahepatic injection
KEGG: Kyoto Encyclopedia of Genes and Genomes
mpPHH: mouse-passaged primary human hepatocytes
PC: precore
pcRNA: precore RNA
pgRNA: pregenomic RNA
PSR: picrosirirus red
rcccDNA: recombinant covalently closed circular DNA
RNA: ribonucleic acid
SILAC: stable isotope labeling by amino acids in cell culture
TCGA: The Cancer Genome Atlas
WT: wild-type.

