## supplementary materials for "The HBV basal core promoter mutation confers a replicative advantage and transcriptionally reprograms hepatocytes toward HCC subtypes"

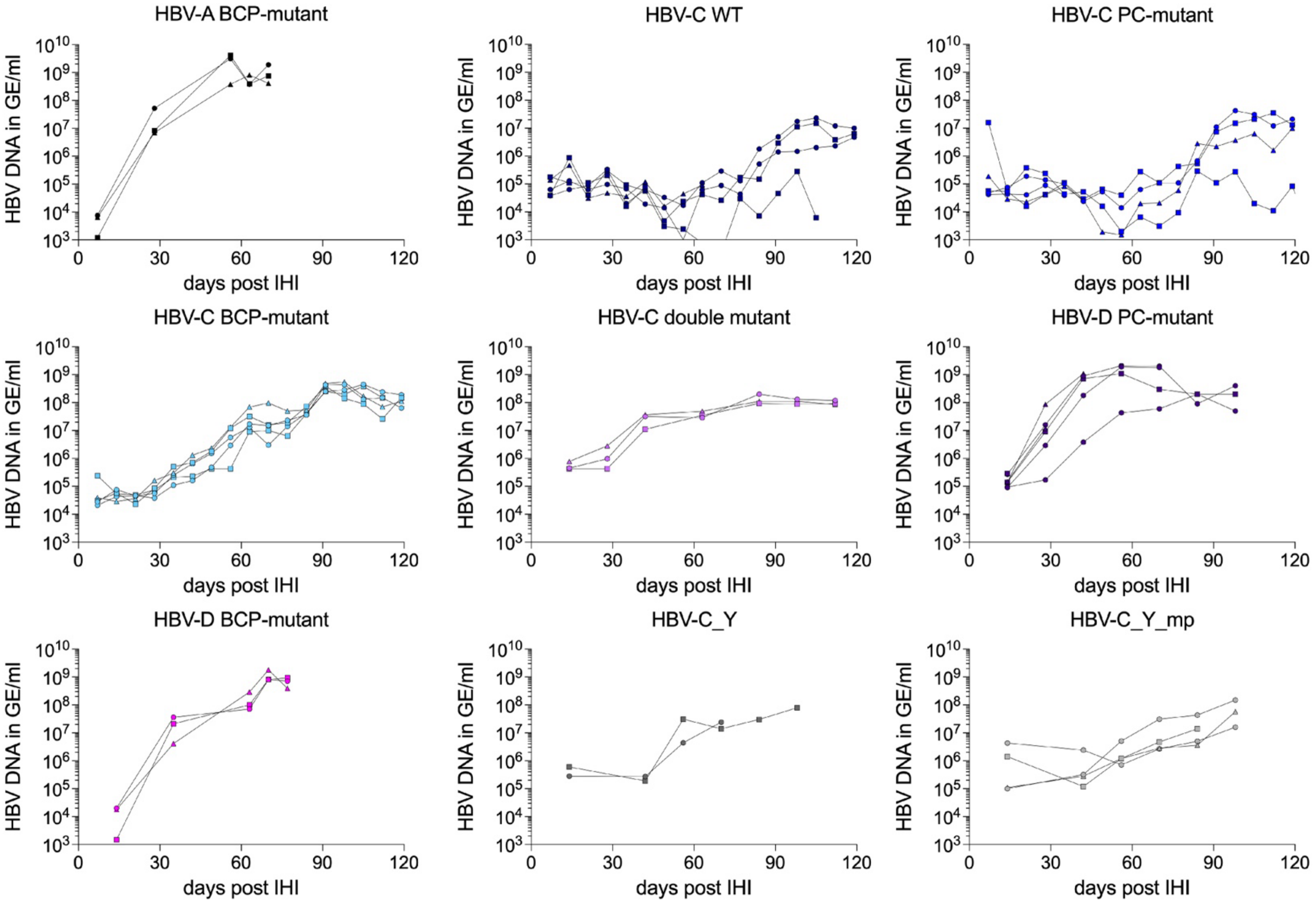

**Figure S1: Successful launch of HBV strains and isogenic variants via IHI rcccDNA across genotypes A–D and patient consensus sequences.**

Serum HBV viremia (HBV DNA in GE/ml) following IHI of rcccDNA in huFNRG mice is shown for nine independent HBV launches of WT strains and BCP- and PC-mutants across genotypes A–D. HBV_C_Y and HBV_CY_mp are patient-derived consensus sequences. *GE = genome equivalents, IHI = intrahepatic injection, rcccDNA = recombinant cccDNA.*

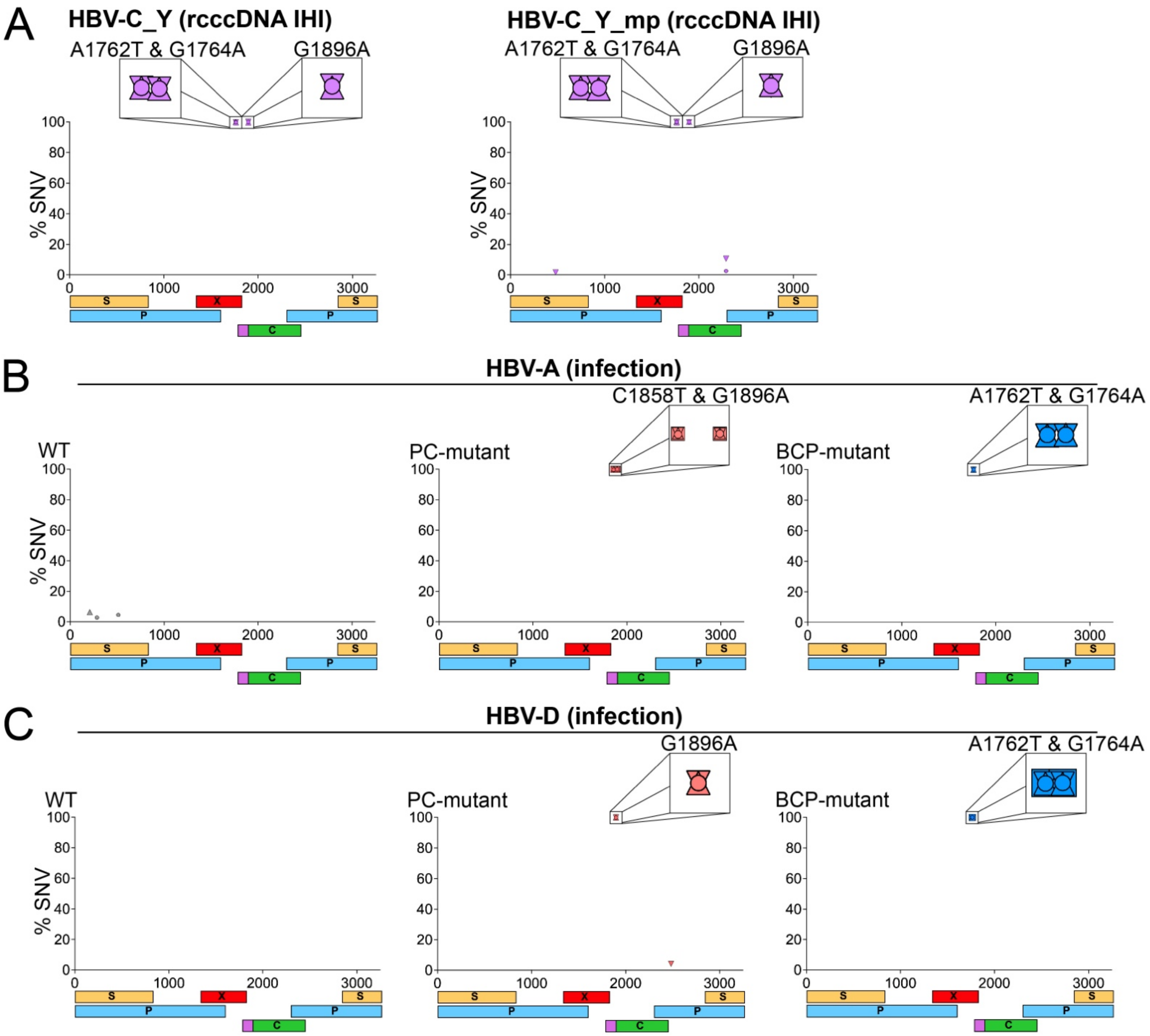

**Figure S2: Viral sequencing of IHI rcccDNA launches and infection experiments.**

**(A)** Sequencing analysis of HBV DNA isolated from serum of huFNRG mice following HBV rcccDNA IHI. Each data point represents an SNV, with each mouse represented by a different symbol (n= 3). The x-axis represents the position on the HBV genome, with the EcoRI restriction site in the S open reading frame (ORF) serving as reference starting point. S = S-antigen ORF, P = polymerase ORF, X = X-protein ORF, C = core ORF (the pink extension represents the ORF unique to the precore protein). **(B)** Sequencing analysis of HBV DNA isolated from serum of huFNRG mice following HBV-A infection. Each data point represents an SNV, with each mouse represented by a different symbol (n= 3-5). **(C)**

Sequencing analysis of HBV DNA isolated from serum of huFNRG mice following HBV-D infection. Each data point represents an SNV, with each mouse represented by a different symbol (n= 3-4). *huFNRG mice = humanized $Fah^{-/-}$ NOD $Rag1^{-/-}$ $Il2rg^{null}$ mice, IHI = intrahepatic injection, rcccDNA = recombinant circular covalently close DNA, SNV = single nucleotide variation.*

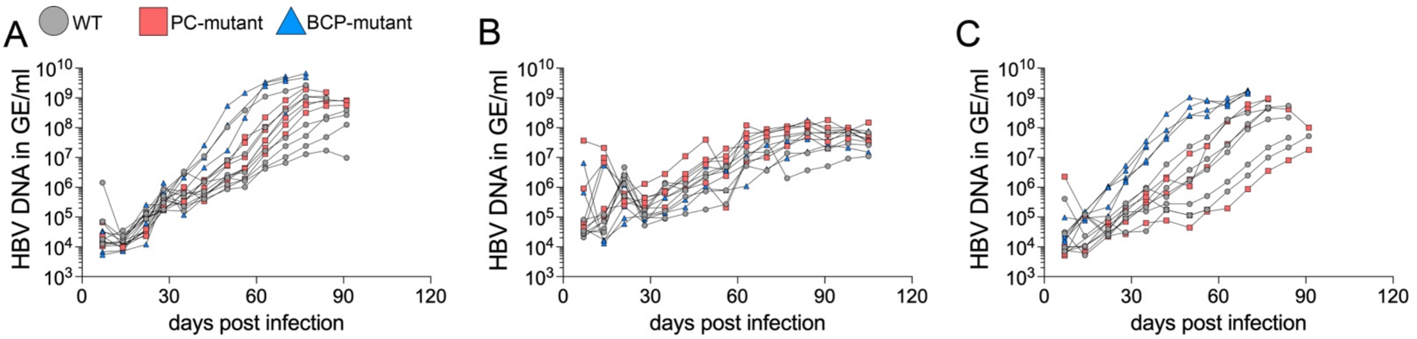

**Figure S3: Hepatitis B infection with isogenic variants in HBV-A, HBV-C, and HBV-D: single mouse viremia.**

*Related to **Fig. 2A**.* Viremia data from individual mice are presented. **(A)** HBV-A infection, **(B)** HBV-C infection, and **(C)** HBV-D infection. Mice were sacrificed at their individual peak plateau viremia level.

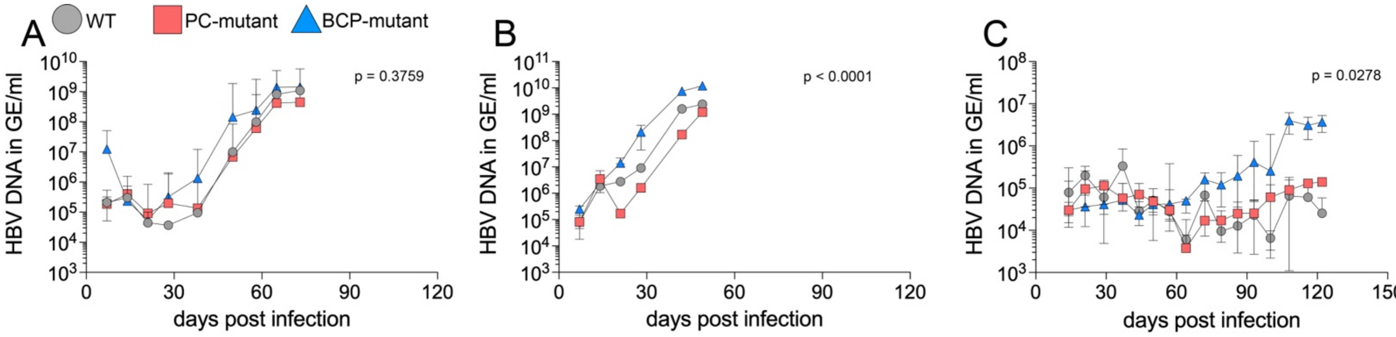

**Figure S4: BCP-mutant infection consistently accelerates viremia kinetics in HBV-D infection.** Independent infection experiments were performed to validate the observed differences in viral kinetics between WT, PC-, and BCP-mutant infection in HBV-D. HBV infections in **(A)** and **(B)** were performed in huFNRG mice engrafted with the same donor as in **Figure 2,** but in independent cohorts. Infections in **(C)** were performed in huFNRG mice engrafted with a different donor, resulting in lower human hepatocyte engraftment and serum human albumin levels (WT cohort: 1.66 mg/ml, range 0.26 - 6.51; PC-mutant cohort: 1.34 mg/ml, range 0.27 - 9.38; BCP-mutant cohort: 1.34 mg/ml, range 0.32 - 6.74). Data are shown as median values per group with interquartile range. Statistical analysis was performed using a linear mixed-effects model fitted by restricted maximum likelihood (REML), including fixed effects for *group*, *time*, and *group* × *time* interaction and a random effect for subject. **(A)** *group*: p = 0.0420; *time*: p = 0.1466; ***group* × *time*: p = 0.3759**, **(B)** *group*: p < 0.0001; *time*: p < 0.0001; ***group* × *time*: p < 0.0001**, **(C)** *group*: p = 0.0481; *time*: p = 0.0003; ***group* × *time*: p = 0.0278**. *GE = genome equivalents, huFNRG mice = humanized Fah$^{-/-}$ NOD Rag1$^{-/-}$ Il2rg$^{null}$ mice*.

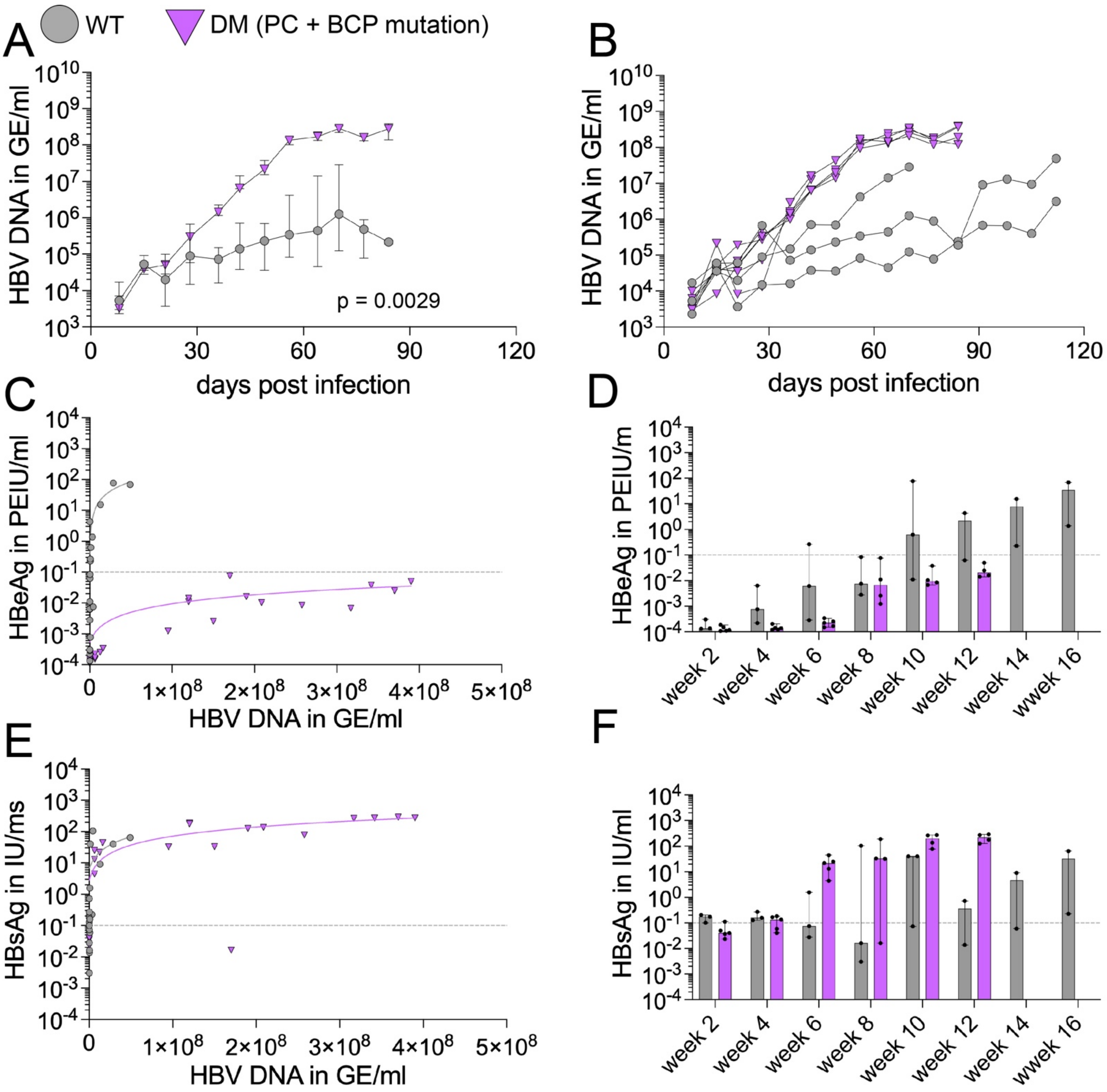

**Figure S5: The double-mutant variant carrying both the PC- and BCP-mutation accelerates viral kinetics in HBV-C.**

Highly humanized huFNRG mice (serum human albumin ~10 mg/mL) were infected with $10^5$ GE via retroorbital injection with either WT (n=3) or double-mutant (DM) (n=6) HBV-C. Serum was collected weekly. **(A)** Grouped serum HBV viremia over time. Data are shown as median with interquartile range. Statistical analysis was performed using a linear mixed-effects model fitted by restricted maximum likelihood (REML), including fixed effects for *group*, *time*, and *group* × *time* interaction and a random effect for subject. (*time*: $p = 0.0023$; *group*: $p < 0.0001$; ***group × time*: p = 0.0029**). **(B)** Serum viremia from individual mice. **(C)** Serum HBeAg levels, measured by chemiluminescence immunoassay, plotted against serum HBV viremia. Each symbol represents a single time point and animal. **(D)** Serum HBeAg levels over time.

**(E)** Serum HBsAg levels, measured by chemiluminescence immunoassay, plotted against serum HBV viremia. Each symbol represents a single time point and animal. **(F)** Serum HBsAg levels over time. *GE = genome equivalent, huFNRG mice = humanized* $Fah^{-/-}$ *NOD* $Rag1^{-/-}$ $Il2rg^{null}$ *mice, IU = international units, LOQ = lower limit of quantification, PEIU = Paul-Ehrlich-Institute units*.

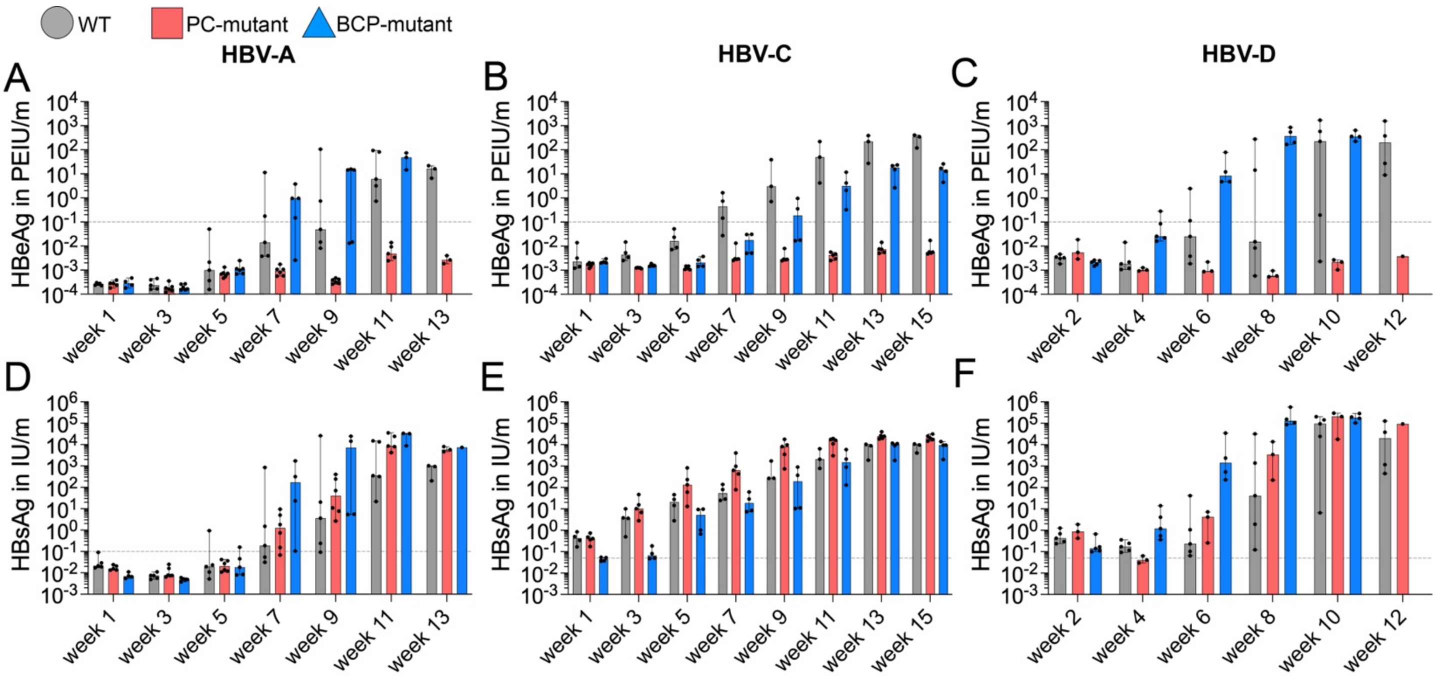

**Figure S6: Time course of serum HBeAg and HBsAg levels in HBV-A, HBV-C, and HBV-D infection.**

*Related to* ***Fig. 2B*** *and* ***2C***. **(A)** Serum HBeAg levels over time. **(B)** Serum HBsAg levels over time. *IU = international units, LOQ = lower limit of quantification, PEIU = Paul-Ehrlich-Institute units*.

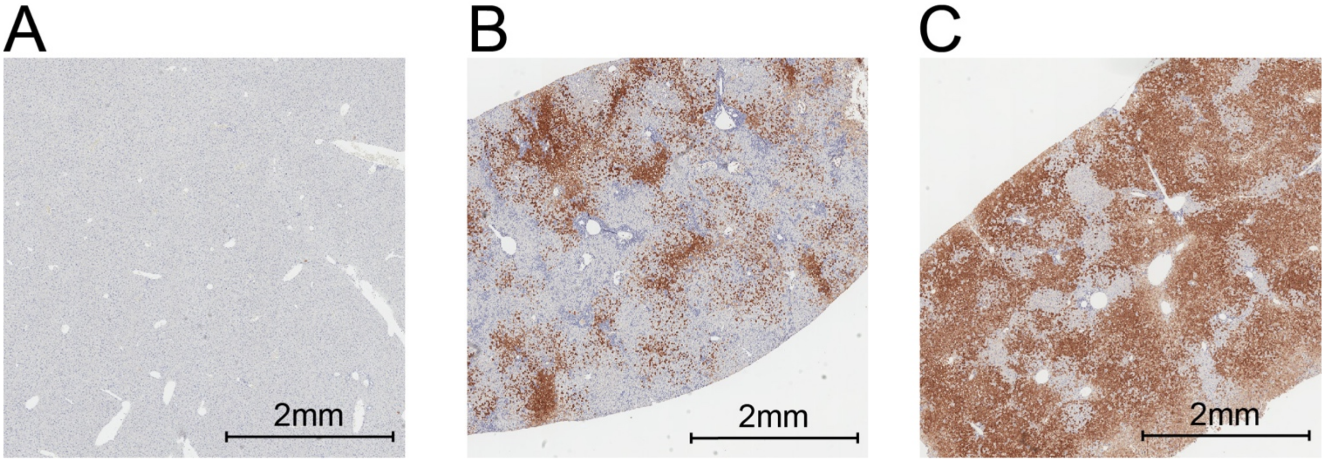

**Figure S7: The human graft remains stable huFNRG mice with HBV infection.**

Shown are representative images of FAH (absent in murine hepatocytes of FNRG mice) in **(A)** non-humanized, **(B)** lowly humanized (1mg/ml serum human albumin) or **(C)** highly humanized (10 mg/ml serum human albumin) huFNRG mice. FAH = fumarylacetoacetate hydrolase, *huFNRG mice* = *humanized* $Fah^{-/-}$ *NOD* $Rag1^{-/-}$ $Il2rg^{null}$ *mice*.

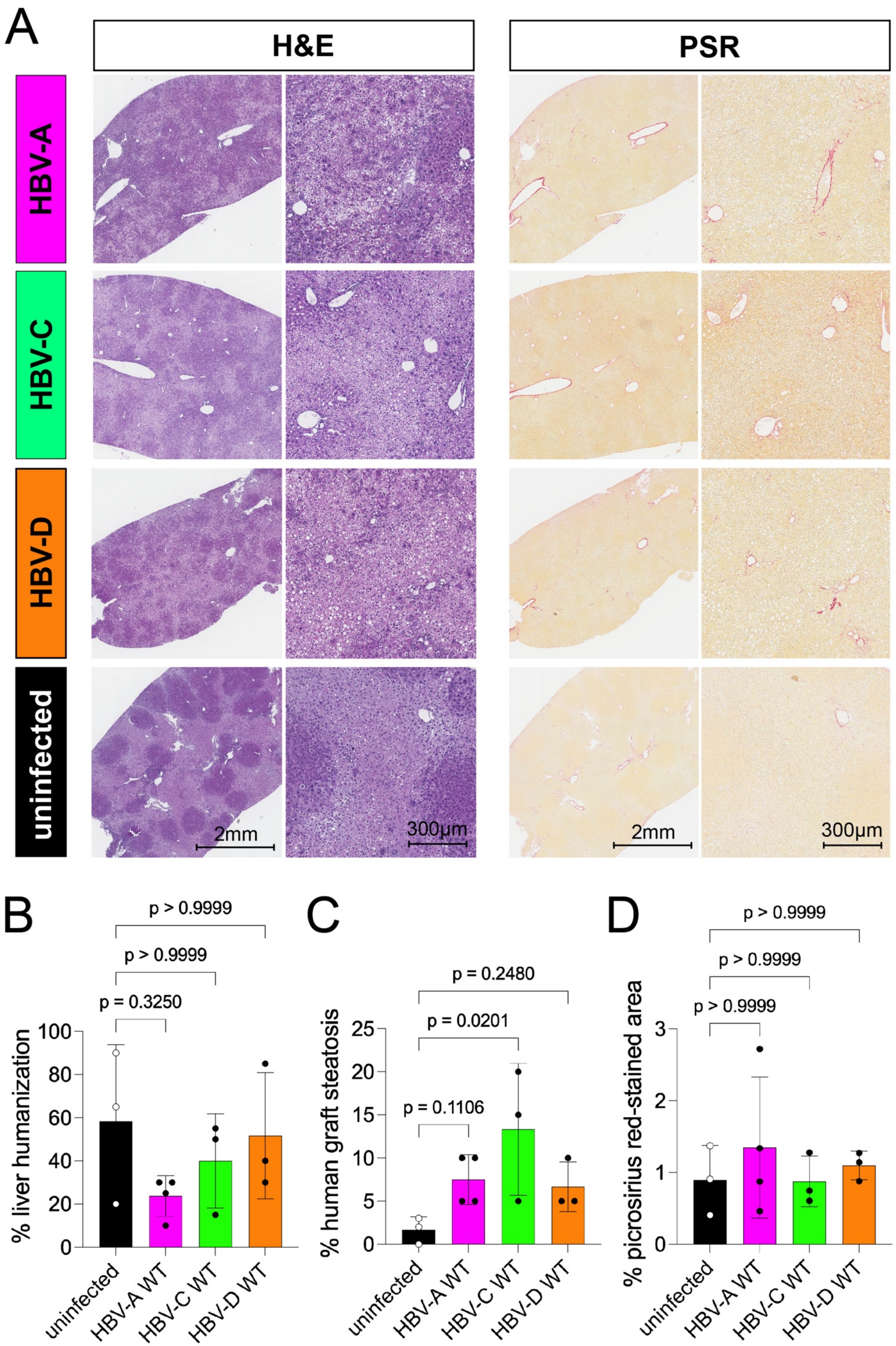

A
H&E
PSR
HBV-A
HBV-C
HBV-D
uninfected
2mm
300µm
2mm
300µm
B
p > 0.9999
p > 0.9999
p = 0.3250
% liver humanization
uninfected
HBV-A WT
HBV-C WT
HBV-D WT
C
p = 0.2480
p = 0.0201
p = 0.1106
% human graft steatosis
uninfected
HBV-A WT
HBV-C WT
HBV-D WT
D
p > 0.9999
p > 0.9999
p > 0.9999
% picrosirius red-stained area
uninfected
HBV-A WT
HBV-C WT
HBV-D WT

**Figure S8: Liver histology in HBV WT infection.**

**(A)** Representative H&E and PSR staining of liver sections from huFNRG mice infected with WT HBV-A, HBV-C, HBV-D, and uninfected controls. **(B)** Percentage of human hepatocytes among all hepatocytes. Statistics: Kruskal-Wallis test with Dunn's multiple comparisons test. **(C)** Percentage of steatosis in the human graft. Statistics: Kruskal-Wallis test with Dunn's multiple comparisons test. **(D)** Picrosirius red-stained area. Statistics: Kruskal-Wallis test with Dunn's multiple comparisons test. *huFNRG mice = humanized Fah$^{-/-}$ NOD Rag1$^{-/-}$ Il2rg$^{null}$ mice*, *H&E = hematoxylin and eosin, PSR = picrosirius red stain*.

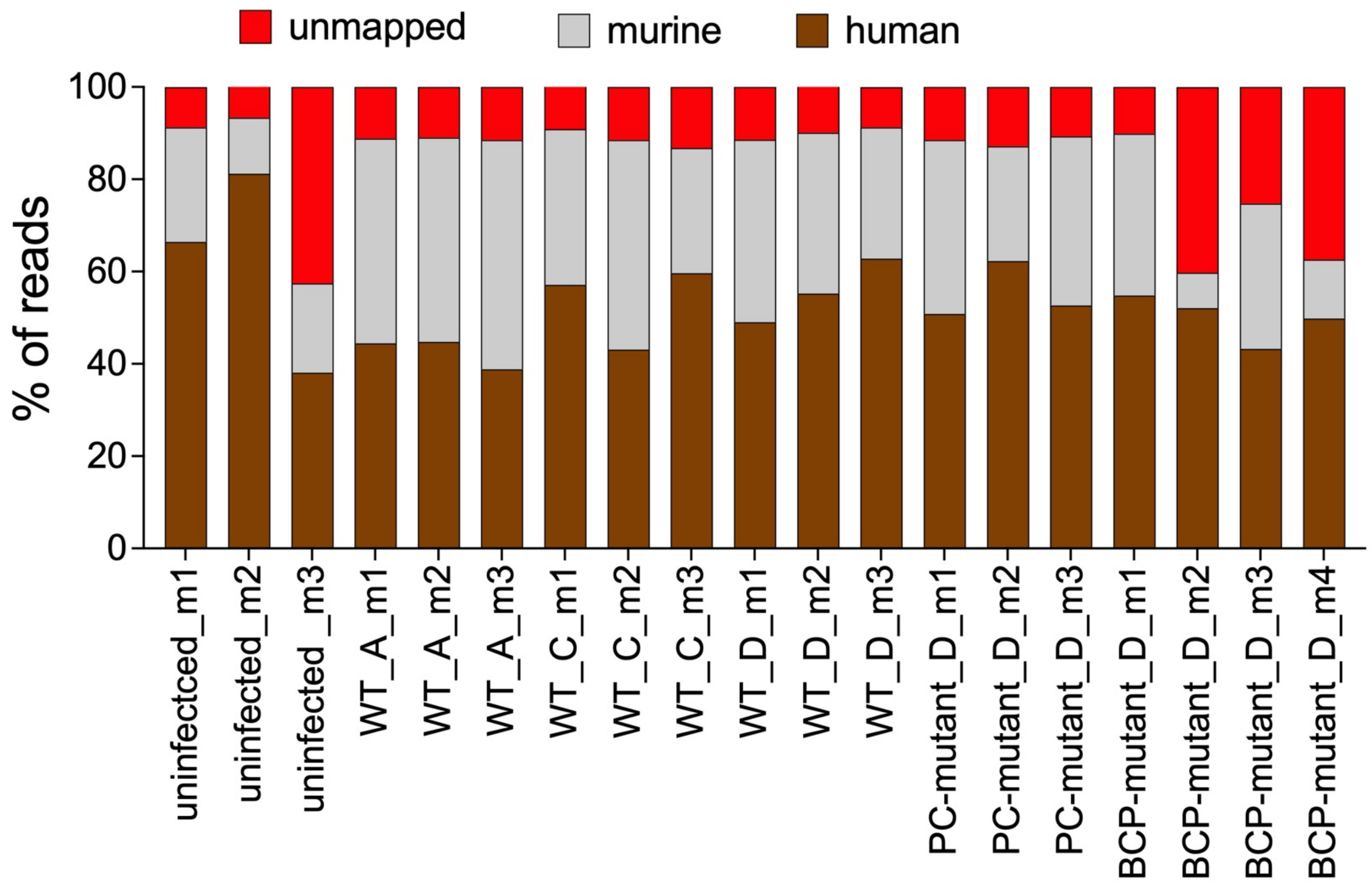

**Figure S9: Proportion of human-, murine-, and ambiguously mapped reads in huFNRG-derived RNA.**

*Related to **Fig. 3**:* Stacked bar plots show the percentage of sequencing reads assigned to the human (brown) or murine genome (grey) or unmapped/ambiguous reads (red) for each sample. Uninfected controls and HBV-infected human hepatocyte chimeric mice (HBV-A WT, HBV-C WT, HBV-D WT, HBV-D BCP-mutant, and HBV-D PC-mutant) are shown. *huFNRG mice = humanized Fah$^{-/-}$ NOD Rag1$^{-/-}$ Il2rg$^{null}$ mice*.

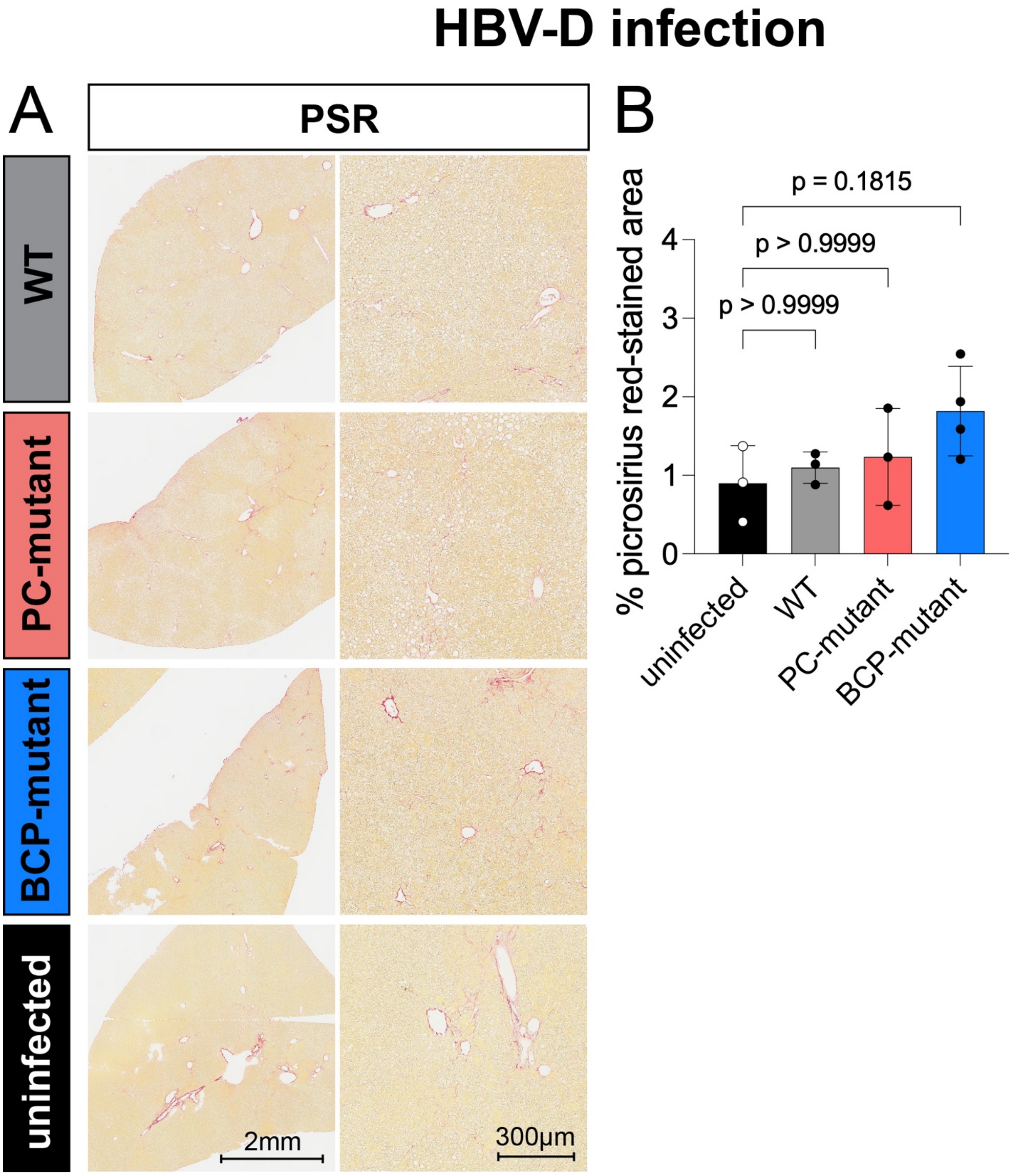

**Figure S10: Picrosirius red-staining in HBV-D infection**

*Related to **Fig. 4**:* **(A)** PSR staining of liver sections from huFNRG mice infected with HBV-D WT, BCP-mutant, PC-mutant, or uninfected controls. **(B)** Picrosirius red-stained area. Statistics: Kruskal-Wallis test with Dunn's multiple comparisons test. *huFNRG mice = humanized Fah$^{-/-}$ NOD Rag1$^{-/-}$ Il2rg$^{null}$ mice, PSR = picrosirius red stain*.

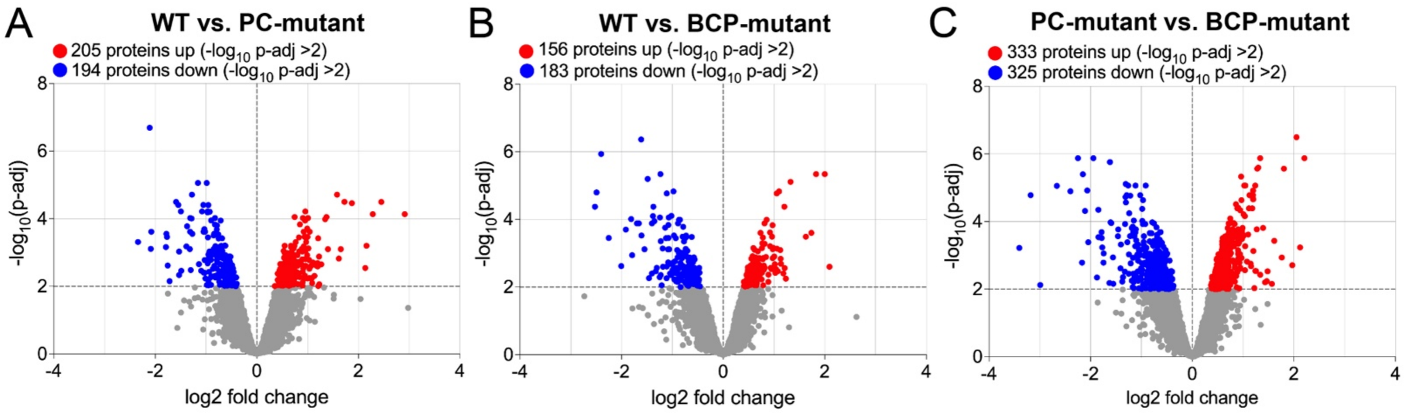

**Figure S11: Volcano plots of differentially expressed proteins identified by SILAC-based proteomics in pre-infected mpPHHs.**

*Related to **Fig. 5**:* Volcano plots show pairwise comparisons of protein abundance (heavy amino acid-labeled) between HBV variants: **(A)** WT vs. PC-mutant, **(B)** WT vs. BCP-mutant, and **(C)** PC- vs. BCP-mutant infection. *mpPHHs = mouse-passaged primary human hepatocytes.*

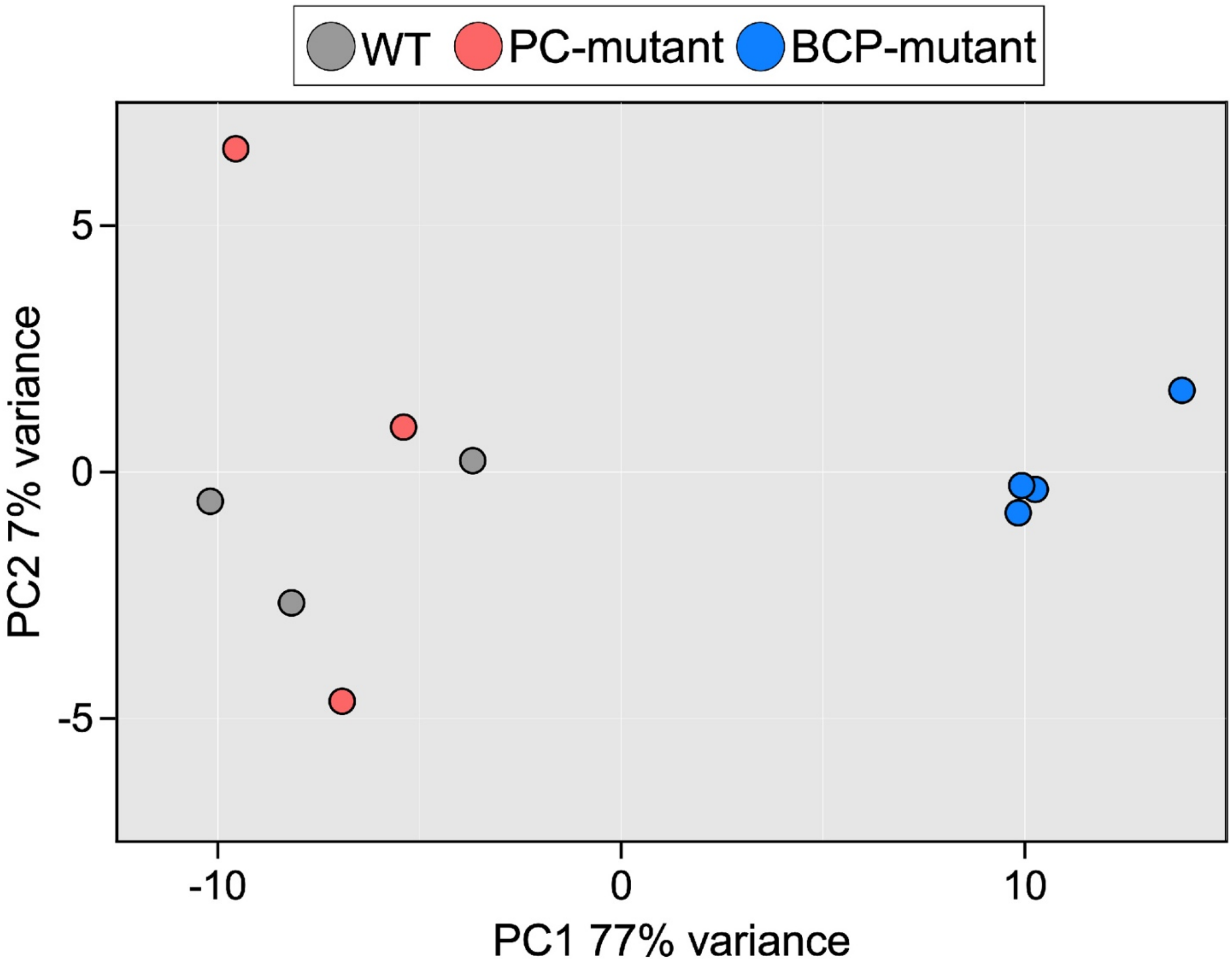

**Figure S12: Principal component analysis of RNA-seq data from bulk liver tissue of huFNRG mice infected with HBV-D WT, BCP-mutant, or PC-mutant.**

*Related to* ***Fig. 6****:* Principal component analysis based on the 1,000 most variable genes between HBV-D WT and BCP-mutant infected huFNRG mice.

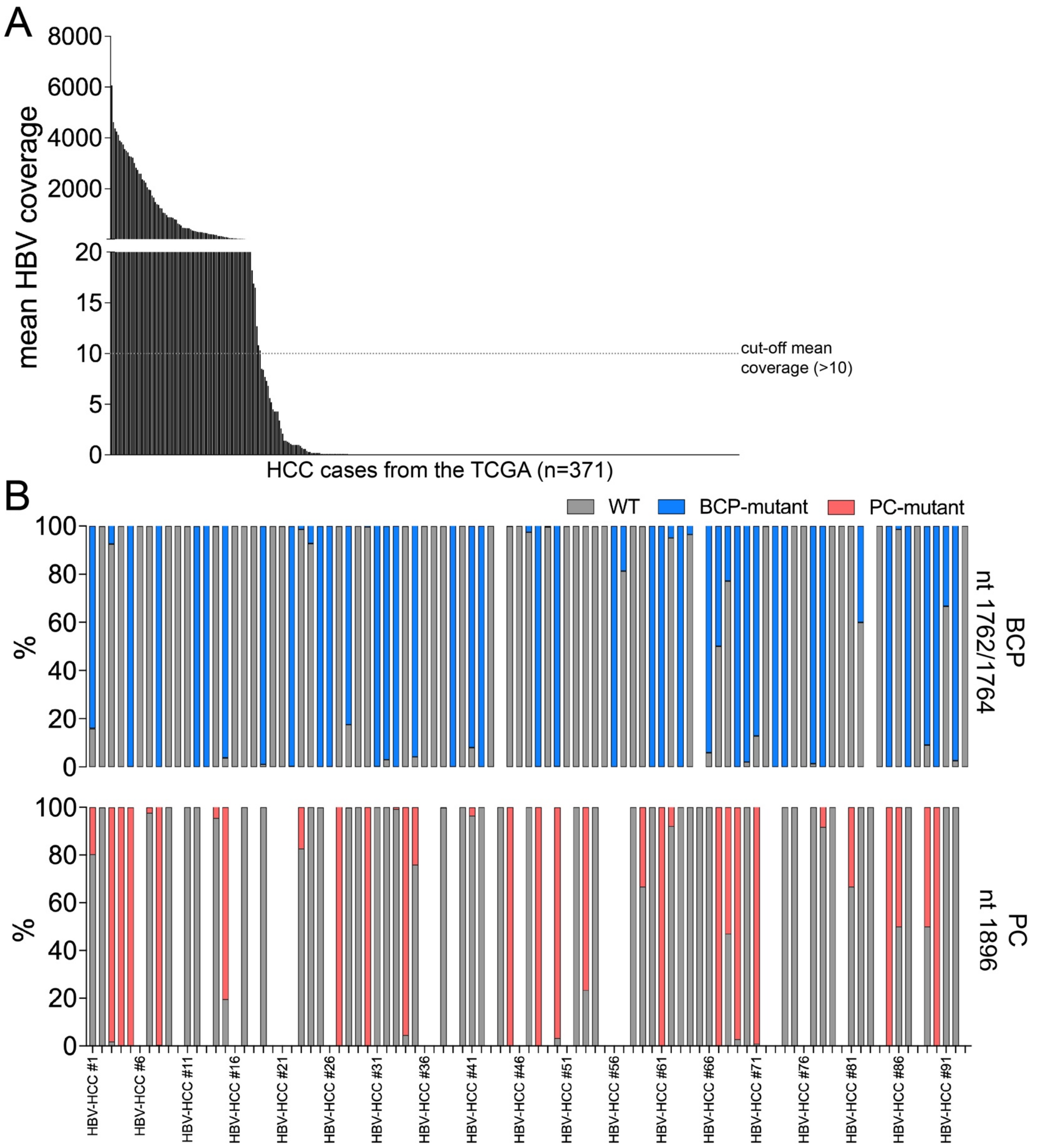

**Figure S13: HBV sequencing coverage and relative variant abundance in human HCC.** *Related to **Fig. 7***: All HCC cases in the TCGA were identified, and available RNA-seq data were accessed to identify HBV-derived RNAs. **(A)** Mean HBV coverage across all HCC cases (n = 371) from

TCGA. **(B)** Relative abundance of BCP- and PC-mutations across all HCC samples with detectable HBV RNA (HBV-HCCs) and a mean HBV coverage > 10. Cases are ordered by mean coverage as in **(A)**. *HCC = hepatocellular carcinoma, TCGA = The Cancer Genome Atlas.*

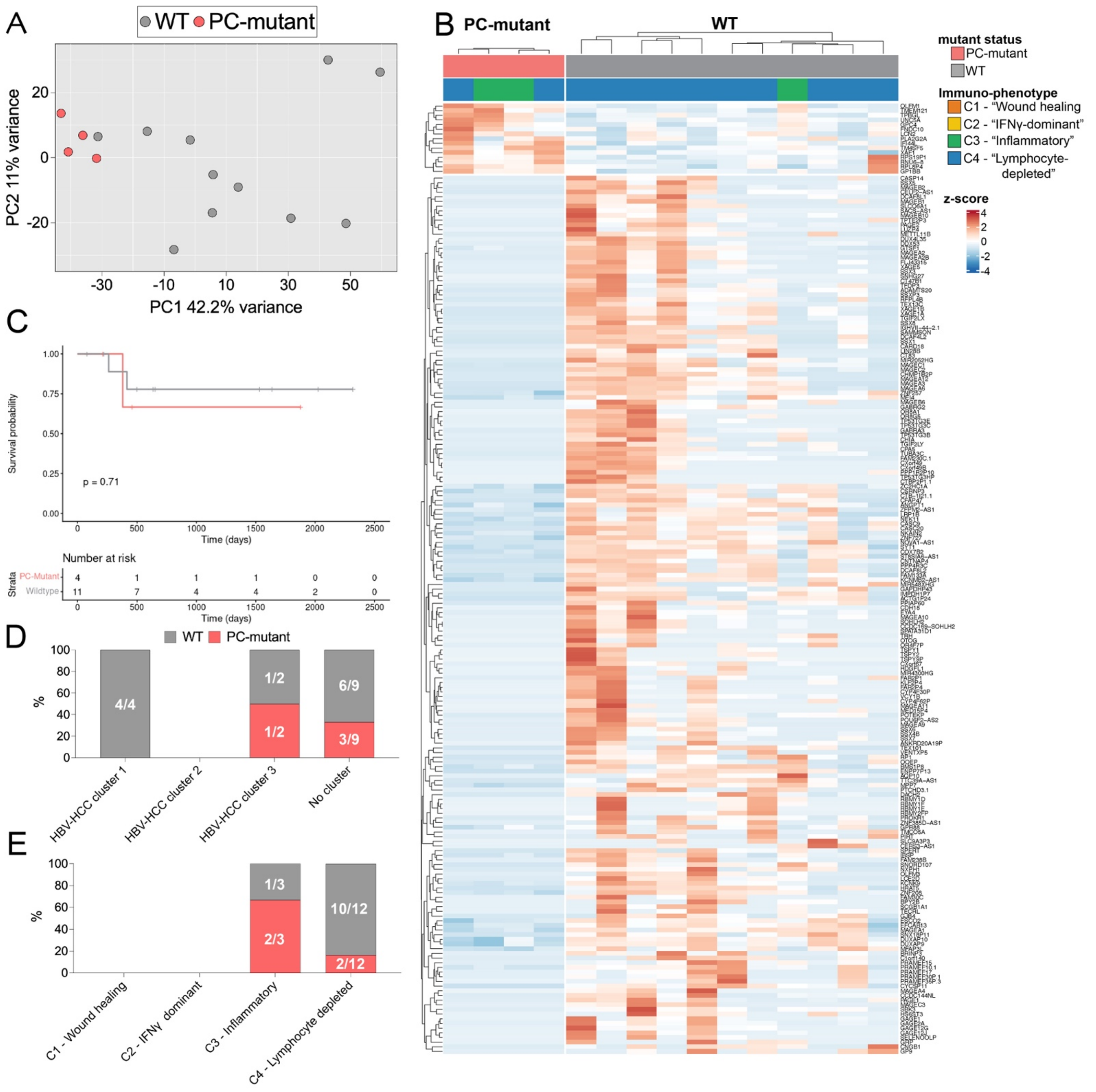

**Figure S14: Comparison of HBV-associated HCCs from the TCGA dataset: PC-mutant vs. WT.** *Related to **Fig. 7***: HBV-HCC cases were classified as PC-mutant or WT if ≥ 97.5% of reads were PC-mutant or WT, respectively, at the PC locus nt1896, and samples with > 0 BCP-mutant reads, or coverage < 10 at the BCP loci nt1762/1764, were excluded ("pure" HBV-HCC samples). **(A)** Principal component analysis HBV-HCCs with WT or "pure" PC-mutant infection based on the 500 most variable genes. **(B)** Heatmap of differentially expressed genes between WT- and PC-mutant associated HCCs

($p$-adj $< 0.001$, log2fc $> |2|$). **(C)** Kaplan-Meier survival analysis comparing WT- and PC-mutant associated HCCs. ($p = 0.71$, long-rank test). **(D and E)** Association of HBV WT- and PC-mutant associated HCCs with the HBV-HCC clusters from **(D)** Tian et al. or **(E)** immune subtypes. *fc = fold change, HCC = hepatocellular carcinoma, TCGA = The Cancer Genome Atlas.*

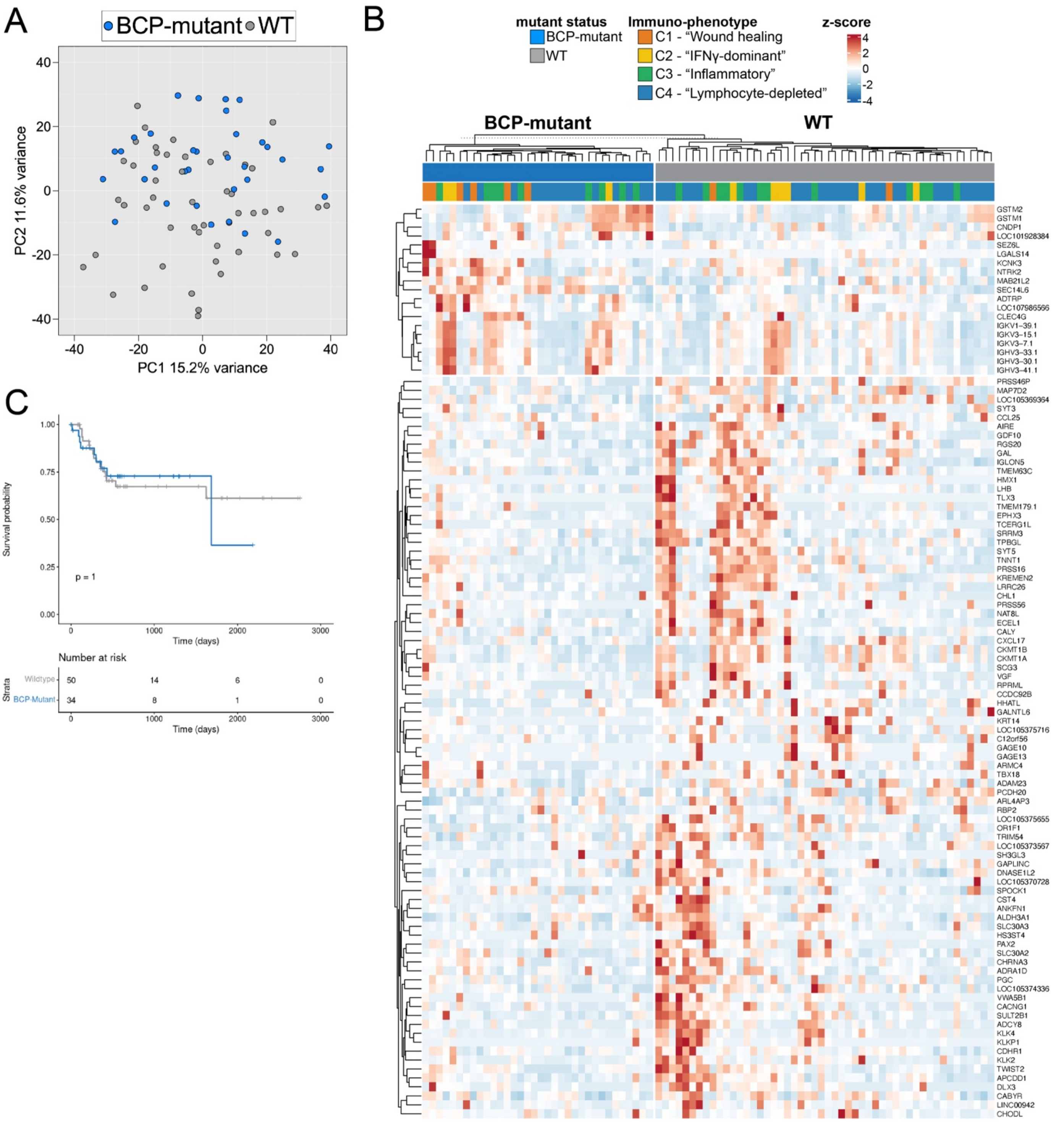

**Figure S15: Comparison of HBV-associated HCCs from the TCGA dataset: BCP-mutant vs. WT ("majority call").**

*Related to* ***Fig. 7***: HBV-HCC cases were classified as BCP-mutant or WT based on the majority of respective reads at position nt1762/1764 (≥ 50%), and samples with PC-mutation ≥ 50% were excluded ("majority call"). **(A)** Principal component analysis HBV-HCCs with WT or BCP-mutant infection based on the 500 most variable genes. **(B)** Heatmap of differentially expressed genes between WT- and BCP -mutant associated HCCs (p-adj < 0.05, log2fc > │2│). **(C)** Kaplan-Meier survival analysis comparing WT- and BCP-mutant associated HCCs. (p = 1, long-rank test). *fc = fold change, HCC = hepatocellular carcinoma, TCGA = The Cancer Genome Atlas.*

**Supplementary tables:**

**Table S1: HBV qPCR primer list**

| **isolate** | **fwd primer ID** | **fwd primer sequence** | **rev primer ID** | **rev primer sequence** | Taqman probe |
|---|---|---|---|---|---|
| **HBV-A** | RU-O-19375 | CCGTCTGTGCCTTCTCATCTG | RU-O-19376 | AGTCCAAGAGTCCTCTTATGTAAGACCTT | 5- /56-FAM/CCG TGT GCA /ZEN/CTT CGCTTC ACCTCT GC/3IABkFQ/ -3 |
| **HBV-B** | RU-O-33045 | CCGTCTGTGCCTTCTCRTCTR | RU-O-32025 | AGTCCAAGAGTCCKCTTATGYAAGACCTT | 5- /56-FAM/CCG TGT GCA /ZEN/CTT CGCTTC ACCTCT GC/3IABkFQ/ -3 |
| **HBV-C** | RU-O-32024 | CCGTCTGTGCCTTCTCRTCTG | RU-O-32025 | AGTCCAAGAGTCCKCTTATGYAAGACCTT | 5- /56-FAM/CCG TGT GCA /ZEN/CTT CGCTTC ACCTCT GC/3IABkFQ/ -3 |
| **HBV-D** | RU-O-19375 | CCGTCTGTGCCTTCTCATCTG | RU-O-19376 | AGTCCAAGAGTCCTCTTATGTAAGACCTT | 5- /56-FAM/CCG TGT GCA /ZEN/CTT CGCTTC ACCTCT GC/3IABkFQ/ -3 |
| **HBV-E** | RU-O-32024 | CCGTCTGTGCCTTCTCRTCTG | RU-O-32025 | AGTCCAAGAGTCCKCTTATGYAAGACCTT | 5- /56-FAM/CCG TGT GCA /ZEN/CTT CGCTTC ACCTCT GC/3IABkFQ/ -3 |
| **HBV-F** | RU-O-32024 | CCGTCTGTGCCTTCTCRTCTG | RU-O-32025 | AGTCCAAGAGTCCKCTTATGYAAGACCTT | 5- /56-FAM/CCG TGT GCA /ZEN/CTT CGCTTC ACCTCT GC/3IABkFQ/ -3 |
| **HBV-CY** | RU-O-32024 | CCGTCTGTGCCTTCTCRTCTG | RU-O-32025 | AGTCCAAGAGTCCKCTTATGYAAGACCTT | 5- /56-FAM/CCG TGT GCA /ZEN/CTT CGCTTC ACCTCT GC/3IABkFQ/ -3 |

**Table S2: HBV sequencing primer list**

| isolate | fwd primer ID | fwd primer sequence | rev primer ID | rev primer sequence |
|---|---|---|---|---|
| **HBV-A** | RU-O-32152 | GCAACTTTTTCACCTCTGCCTAATCATC | RU-O-34056 | CATGGTGCTGGTGCGCAG |
| **HBV-B** | RU-O-32152 | GCAACTTTTTCACCTCTGCCTAATCATC | RU-O-34063 | CATGGTGCTGGTGAACAC |
| **HBV-C** | RU-O-32152 | GCAACTTTTTCACCTCTGCCTAATCATC | RU-O-34070 | CATGGTGCTGGTGAACAG |
| **HBV-D** | RU-O-32152 | GCAACTTTTTCACCTCTGCCTAATCATC | RU-O-34056 | CATGGTGCTGGTGCGCAG |
| **HBV-E** | RU-O-32152 | GCAACTTTTTCACCTCTGCCTAATCATC | RU-O-34056 | CATGGTGCTGGTGCGCAG |
| **HBV-F** | RU-O-32152 | GCAACTTTTTCACCTCTGCCTAATCATC | RU-O-34070 | CATGGTGCTGGTGAACAG |
| **HBV-CY** | RU-O-32152 | GCAACTTTTTCACCTCTGCCTAATCATC | RU-O-34070 | CATGGTGCTGGTGAACAG |

**Table S3: Serum HBeAg levels in huFNRG mice**

| **Group ID** | **n** | **Median $\Delta\log_{10}$** | **IQR (Q1–Q3)** | **Fold change (WT / BCP)** | **p value (Mann–Whitney U, two-sided)** |
|---|---|---|---|---|---|
| **HBV-A WT** | 5 | –7.2389 | –7.2889 to –7.2155 | 6.62× higher than BCP | — |
| **HBV-A BCP mutant** | 4 | –8.0598 | –8.0839 to –7.9977 | reference (1×) | 0.0159 |
| **Group ID** | **n** | **Median $\Delta\log_{10}$** | **IQR (Q1–Q3)** | **Fold change (WT / BCP)** | **p value (Mann–Whitney U, two-sided)** |
| **HBV-C WT** | 3 | –5.259 | –5.864 to –5.065 | 32.82× higher than BCP | — |
| **HBV-C BCP mutant** | 4 | –6.775 | –6.934 to –6.614 | reference (1×) | 0.0571 |
| **Group ID** | **n** | **Median $\Delta\log_{10}$** | **IQR (Q1–Q3)** | **Fold change (WT / BCP)** | **p value (Mann–Whitney U, two-sided)** |
| **HBV-D WT** | 5 | –5.7694 | –6.2245 to –5.6589 | 7.22× higher than BCP | — |
| **HBV-D BCP mutant** | 4 | –6.6279 | –6.7213 to –6.5549 | reference (1×) | 0.0159 |
| **Group (WT)** | **n** | **Median $\Delta\log_{10}$** | **IQR (Q1–Q3)** | | |
| **HBV-A WT** | 5 | –7.2389 | –7.2889 to –7.2155 | | |
| **HBV-C WT** | 3 | –5.2590 | –5.8641 to –5.0651 | | |
| **HBV-D WT** | 5 | –5.7694 | –6.2245 to –5.6589 | | |
| **A vs C (fold)** | **A vs C (p)** | **A vs D (fold)** | **A vs D (p)** | **C vs D (fold)** | **C vs D (p)** |
| 0.010× | 0.0357 | 0.034× | 0.0079 | — | — |
| 95.5× | 0.0357 | — | — | 3.24× | 0.571 |
| — | — | 29.5× | 0.0079 | 0.31× | 0.571 |

### Table S4: Serum HBsAg levels in huFNRG mice

| HBV-A | | | | | |
|---|---|---|---|---|---|
| **Group** | **n** | **Median $\Delta\log_{10}$** | **IQR (Q1–Q3)** | | |
| **WT** | 19 | –5.2893 | –6.4418 to –5.0579 | | |
| **PC** | 18 | –5.0680 | –6.1425 to –4.8236 | | |
| **BCP** | 15 | –5.4519 | –5.7328 to –5.1863 | | |
| **WT vs PC (fold)** | **WT vs PC (p)** | **WT vs BCP (fold)** | **WT vs BCP (p)** | **PC vs BCP (fold)** | **PC vs BCP (p)** |
| 0.6008× | 0.2945 | 1.4543× | 0.7287 | — | — |
| 1.6644× | 0.2945 | — | — | 2.4205× | 0.1335 |
| — | — | 0.6876× | 0.7287 | 0.4131× | 0.1335 |
| **HBV-C** | | | | | |
| **Group** | **n** | **Median $\Delta\log_{10}$** | **IQR (Q1–Q3)** | | |
| **WT** | 28 | –4.3578 | –4.6788 to –3.8187 | | |
| **PC** | 40 | –3.8831 | –4.2198 to –3.6210 | | |
| **BCP** | 29 | –4.6842 | –5.1737 to –4.1535 | | |
| **WT vs PC (fold)** | **WT vs PC (p)** | **WT vs BCP (fold)** | **WT vs BCP (p)** | **PC vs BCP (fold)** | **PC vs BCP (p)** |
| 0.34× | 0.0718 | 2.12× | 0.0487 | — | — |
| 2.98× | 0.0718 | — | — | 6.33× | $4.7\times10^{-4}$ |
| — | — | 0.47× | 0.0487 | 0.16× | $4.7\times10^{-4}$ |
| **HBV-D** | | | | | |
| **Group** | **n** | **Median $\Delta\log_{10}$** | **IQR (Q1–Q3)** | | |
| **WT** | 34 | –4.7428 | –5.8355 to –3.6852 | | |
| **PC** | 17 | –3.8506 | –5.4316 to –3.6495 | | |
| **BCP** | 22 | –5.1210 | –5.8950 to –3.9136 | | |
| **WT vs PC (fold)** | **WT vs PC (p)** | **WT vs BCP (fold)** | **WT vs BCP (p)** | **PC vs BCP (fold)** | **PC vs BCP (p)** |
| 0.13× | 0.353 | 2.39× | 0.551 | — | — |
| 7.80× | 0.353 | — | — | 18.64× | 0.098 |
| — | — | 0.42× | 0.551 | 0.054× | 0.098 |
| **Group comparison** | | | | | |
| **group** | **n** | **Median $\Delta\log_{10}$** | **IQR (Q1–Q3)** | | |
| **HBV-A WT** | 19 | –5.2893 | –6.4418 to –5.0579 | | |
| **HBV-C WT** | 28 | –4.3578 | –4.6788 to –3.8187 | | |
| **HBV-D WT** | 34 | –4.7428 | –5.8355 to –3.6852 | | |
| **A vs C (fold)** | **A vs C (p)** | **A vs D (fold)** | **A vs D (p)** | **C vs D (fold)** | **C vs D (p)** |
| 0.12× | 0.0023 | 0.28× | 0.018 | — | — |
| 8.54× | 0.0023 | — | — | 2.43× | 0.041 |
| — | — | 3.52× | 0.018 | 0.41× | 0.041 |

**Table S5: Dysregulated KEGG pathways in SILAC proteomics in HBV-D infected mpPHHs**

| Table S5A: BCP-mutant vs. WT (up) | | | | |
|---|---|---|---|---|
| **Enrichment FDR** | **nGenes** | **Pathway Genes** | **Fold Enrichment** | **Pathway** |
| 2.14E-13 | 20 | 205 | 11.5769039 | Path:hsa05203 Viral carcinogenesis |
| 3.98E-12 | 18 | 190 | 11.241783 | Path:hsa04613 Neutrophil extracellular trap formation |
| 3.98E-12 | 16 | 138 | 13.7580597 | Path:hsa05322 Systemic lupus erythematosus |
| 4.79E-08 | 14 | 188 | 8.83662614 | Path:hsa05034 Alcoholism |
| 6.47E-07 | 14 | 234 | 7.0995116 | Path:hsa05171 Coronavirus disease-COVID-19 |
| 3.76E-06 | 35 | 1556 | 2.66916085 | Path:hsa01100 Metabolic pathways |
| 5.61E-06 | 12 | 202 | 7.04930289 | Path:hsa05169 Epstein-Barr virus infection |
| 0.00010449 | 10 | 181 | 6.55598151 | Path:hsa05168 Herpes simplex virus 1 infection |
| 0.00042758 | 9 | 171 | 6.24543502 | Path:hsa05164 Influenza A |
| 0.00064212 | 6 | 69 | 10.3185448 | Path:hsa05416 Viral myocarditis |
| 0.00132008 | 8 | 159 | 5.97047876 | Path:hsa04514 Cell adhesion molecules |
| 0.00132008 | 6 | 80 | 8.8997449 | Path:hsa04612 Antigen processing and presentation |
| 0.00147825 | 9 | 210 | 5.08556851 | Path:hsa05170 Human immunodeficiency virus 1 infection |
| 0.00159783 | 6 | 86 | 8.27883246 | Path:hsa04610 Complement and coagulation cascades |
| 0.00299587 | 5 | 63 | 9.41771947 | Path:hsa03250 Viral life cycle-HIV-1 |
| 0.00299587 | 7 | 139 | 5.97584789 | Path:hsa05162 Measles |
| 0.00368372 | 8 | 194 | 4.89333053 | Path:hsa05167 Kaposi sarcoma-associated herpesvirus infection |
| 0.00402308 | 4 | 38 | 12.49087 | Path:hsa05330 Allograft rejection |
| 0.00430661 | 7 | 153 | 5.42903828 | Path:hsa01240 Biosynthesis of cofactors |
| 0.00448777 | 5 | 72 | 8.24050454 | Path:hsa04622 RIG-I-like receptor signaling pathway |
| 0.0049155 | 7 | 159 | 5.22416891 | Path:hsa05160 Hepatitis C |
| 0.00532541 | 4 | 43 | 11.0384433 | Path:hsa04940 Type I diabetes mellitus |
| 0.00556661 | 4 | 44 | 10.7875696 | Path:hsa05332 Graft-versus-host disease |
| 0.00654217 | 5 | 82 | 7.23556496 | Path:hsa04623 Cytosolic DNA-sensing pathway |
| 0.00654217 | 8 | 223 | 4.25697813 | Path:hsa05166 Human T-cell leukemia virus 1 infection |
| 0.00741011 | 5 | 85 | 6.98019208 | Path:hsa01232 Nucleotide metabolism |
| 0.0096428 | 4 | 53 | 8.95571814 | Path:hsa05320 Autoimmune thyroid disease |
| 0.01882525 | 9 | 332 | 3.21677526 | Path:hsa05165 Human papillomavirus infection |
| 0.01894295 | 6 | 157 | 4.53490186 | Path:hsa04218 Cellular senescence |
| 0.0249792 | 7 | 224 | 3.70822704 | Path:hsa05163 Human cytomegalovirus infection |
| 0.04134813 | 5 | 131 | 4.52913226 | Path:hsa04650 Natural killer cell mediated cytotoxicity |

| | | | | |
|---|---|---|---|---|
| 0.04254902 | 7 | 250 | 3.32257143 | Path:hsa04144 Endocytosis |
| 0.0454276 | 5 | 136 | 4.36262005 | Path:hsa00190 Oxidative phosphorylation |

**Table S5B: BCP-mutant vs. WT (down)**

| Enrichment FDR | nGenes | Pathway Genes | Fold Enrichment | Pathway |
|---|---|---|---|---|
| 4.05E-07 | 12 | 170 | 10.5918786 | Path:hsa04141 Protein processing in endoplasmic reticulum |
| 1.51E-05 | 11 | 203 | 8.13087558 | Path:hsa05205 Proteoglycans in cancer |
| 2.12E-05 | 9 | 132 | 10.2307918 | Path:hsa04142 Lysosome |
| 2.88E-05 | 11 | 232 | 7.11451613 | Path:hsa04820 Cytoskeleton in muscle cells |
| 5.27E-05 | 10 | 202 | 7.42829767 | Path:hsa04510 Focal adhesion |
| 0.00119592 | 6 | 89 | 10.1158391 | Path:hsa04512 ECM-receptor interaction |
| 0.00121916 | 25 | 1556 | 2.41085496 | Path:hsa01100 Metabolic pathways |
| 0.00121916 | 7 | 138 | 7.6113137 | Path:hsa04915 Estrogen signaling pathway |
| 0.00447031 | 10 | 362 | 4.14507218 | Path:hsa04151 PI3K-Akt signaling pathway |
| 0.00709079 | 3 | 20 | 22.5077419 | Path:hsa00100 Steroid biosynthesis |
| 0.00709079 | 4 | 49 | 12.2491113 | Path:hsa00330 Arginine and proline metabolism |
| 0.01131209 | 6 | 152 | 5.92308998 | Path:hsa04145 Phagosome |
| 0.01197152 | 6 | 156 | 5.77121588 | Path:hsa04148 Efferocytosis |
| 0.01634125 | 7 | 231 | 4.54701857 | Path:hsa04810 Reg. of actin cytoskeleton |
| 0.0225609 | 4 | 75 | 8.00275269 | Path:hsa01230 Biosynthesis of amino acids |
| 0.0225609 | 2 | 9 | 33.3448029 | Path:hsa03266 Virion-Herpesvirus |
| 0.0225609 | 5 | 133 | 5.64103808 | Path:hsa04270 Vascular smooth muscle contraction |
| 0.0225609 | 4 | 80 | 7.50258065 | Path:hsa04612 Antigen processing and presentation |
| 0.0225609 | 5 | 129 | 5.81595399 | Path:hsa04926 Relaxin signaling pathway |
| 0.0225609 | 4 | 78 | 7.69495451 | Path:hsa05100 Bacterial invasion of epithelial cells |
| 0.0225609 | 7 | 248 | 4.23532778 | Path:hsa05132 Salmonella infection |
| 0.0225609 | 8 | 332 | 3.61570152 | Path:hsa05165 Human papillomavirus infection |
| 0.02499111 | 5 | 138 | 5.43665264 | Path:hsa04910 Insulin signaling pathway |
| 0.0295335 | 6 | 210 | 4.28718894 | Path:hsa05170 Human immunodeficiency virus 1 infection |
| 0.0296563 | 5 | 147 | 5.10379636 | Path:hsa04072 Phospholipase D signaling pathway |
| 0.0296563 | 3 | 44 | 10.2307918 | Path:hsa04962 Vasopressin-regulated water reabsorption |
| 0.03174898 | 4 | 93 | 6.45383281 | Path:hsa04657 IL-17 signaling pathway |
| 0.03322186 | 3 | 47 | 9.57776253 | Path:hsa00514 Other types of O-glycan biosynthesis |
| 0.03322186 | 6 | 224 | 4.01923963 | Path:hsa05163 Human cytomegalovirus infection |
| 0.03322186 | 4 | 97 | 6.18769538 | Path:hsa05215 Prostate cancer |

| | | | | |
|---|---|---|---|---|
| 0.03349376 | 4 | 99 | 6.06269143 | Path:hsa04933 AGE-RAGE signaling pathway in diabetic complications |
| 0.03349376 | 4 | 99 | 6.06269143 | Path:hsa05231 Choline metabolism in cancer |
| 0.03610724 | 4 | 102 | 5.88437698 | Path:hsa05146 Amoebiasis |
| 0.03679179 | 6 | 236 | 3.81487151 | Path:hsa04014 Ras signaling pathway |
| 0.03797343 | 5 | 168 | 4.46582181 | Path:hsa04140 Autophagy-animal |
| 0.04651399 | 2 | 19 | 15.7949066 | Path:hsa00531 Glycosaminoglycan degradation |
| 0.04651399 | 4 | 115 | 5.21918654 | Path:hsa01200 Carbon metabolism |
| 0.04651399 | 4 | 115 | 5.21918654 | Path:hsa04724 Glutamatergic synapse |
| 0.04651399 | 5 | 181 | 4.14507218 | Path:hsa05168 Herpes simplex virus 1 infection |
| **Table S5C: PC-mutant vs. WT (up)** | | | | |
| **Enrichment FDR** | **nGenes** | **Pathway Genes** | **Fold Enrichment** | **Pathway** |
| 5.00E-05 | 34 | 1556 | 2.61963003 | Path:hsa01100 Metabolic pathways |
| 0.00220768 | 8 | 136 | 7.05215282 | Path:hsa00190 Oxidative phosphorylation |
| 0.00321704 | 6 | 76 | 9.46473142 | Path:hsa03320 PPAR signaling pathway |
| 0.03555905 | 5 | 85 | 7.05215282 | Path:hsa01232 Nucleotide metabolism |
| 0.03555905 | 5 | 86 | 6.97015104 | Path:hsa04610 Complement and coagulation cascades |
| 0.04219503 | 5 | 93 | 6.44551602 | Path:hsa04520 Adherens junction |
| **Table S5D: PC-mutant vs. WT (down)** | | | | |
| **Enrichment FDR** | **nGenes** | **Pathway Genes** | **Fold Enrichment** | **Pathway** |
| 1.98E-09 | 16 | 185 | 9.76457623 | Path:hsa03040 Spliceosome |
| 3.44E-05 | 9 | 103 | 9.86530304 | Path:hsa03015 mRNA surveillance pathway |
| 0.00050422 | 11 | 232 | 5.35315534 | Path:hsa04820 Cytoskeleton in muscle cells |
| 0.00119017 | 8 | 132 | 6.84260076 | Path:hsa04142 Lysosome |
| 0.00190532 | 7 | 107 | 7.38617185 | Path:hsa03013 Nucleocytoplasmic transport |
| 0.00473835 | 8 | 170 | 5.31307824 | Path:hsa04141 Protein processing in endoplasmic reticulum |
| 0.01037801 | 9 | 246 | 4.13059436 | Path:hsa05131 Shigellosis |
| 0.01985174 | 7 | 169 | 4.676452 | Path:hsa04530 Tight junction |
| 0.02369016 | 8 | 231 | 3.91005758 | Path:hsa04810 Reg. of actin cytoskeleton |
| 0.02369016 | 7 | 181 | 4.36641099 | Path:hsa05168 Herpes simplex virus 1 infection |
| 0.02564974 | 6 | 137 | 4.94465311 | Path:hsa04210 Apoptosis |
| 0.04273059 | 3 | 31 | 10.9260883 | Path:hsa03060 Protein export |

**Table S6: KEGG pathways enriched in HBV-D BCP-mutant infected huFNRG mice (RNAseq)**

| Enrichment FDR | nGenes | Pathway Genes | Fold Enrichment | Pathway |
|---|---|---|---|---|
| 1.51E-21 | 377 | 1556 | 1.63100793 | Path:hsa01100 Metabolic pathways |
| 4.37E-10 | 59 | 158 | 2.51373354 | Path:hsa04110 Cell cycle |
| 4.37E-10 | 139 | 529 | 1.7688192 | Path:hsa05200 Pathways in cancer |
| 2.37E-09 | 23 | 36 | 4.30080399 | Path:hsa03030 DNA replication |
| 4.85E-07 | 55 | 170 | 2.17790074 | Path:hsa04141 Protein processing in endoplasmic reticulum |
| 8.03E-05 | 45 | 149 | 2.0330617 | Path:hsa05226 Gastric cancer |
| 0.00017845 | 48 | 168 | 1.92334091 | Path:hsa05225 Hepatocellular carcinoma |
| 0.0002196 | 74 | 300 | 1.66048432 | Path:hsa04010 MAPK signaling pathway |
| 0.00066137 | 27 | 80 | 2.27194645 | Path:hsa00983 Drug metabolism-other enzymes |
| 0.00066137 | 33 | 107 | 2.07612968 | Path:hsa04350 TGF-beta signaling pathway |
| 0.00066137 | 105 | 478 | 1.47871922 | Path:hsa05022 Pathways of neurodegeneration-multiple diseases |
| 0.00076593 | 19 | 48 | 2.66462856 | Path:hsa03272 Virion-Hepatitis viruses |
| 0.00083479 | 12 | 23 | 3.51218776 | Path:hsa03430 Mismatch repair |
| 0.00087871 | 11 | 20 | 3.70243126 | Path:hsa00100 Steroid biosynthesis |
| 0.00113839 | 21 | 58 | 2.43733719 | Path:hsa00480 Glutathione metabolism |
| 0.00134417 | 31 | 103 | 2.02604358 | Path:hsa04064 NF-kappa B signaling pathway |
| 0.00180956 | 40 | 148 | 1.81937654 | Path:hsa05224 Breast cancer |
| 0.00193805 | 42 | 159 | 1.77818311 | Path:hsa04217 Necroptosis |
| 0.00193805 | 53 | 215 | 1.65944065 | Path:hsa05207 Chemical carcinogenesis-receptor activation |
| 0.0021437 | 26 | 84 | 2.08361932 | Path:hsa04012 ErbB signaling pathway |
| 0.0021437 | 59 | 248 | 1.60149153 | Path:hsa05132 Salmonella infection |
| 0.0021437 | 28 | 93 | 2.02674634 | Path:hsa05222 Small cell lung cancer |
| 0.0021437 | 38 | 141 | 1.81421519 | Path:hsa05418 Fluid shear stress and atherosclerosis |

| | | | | |
|---|---|---|---|---|
| 0.0022984 | 37 | 137 | 1.8180485<br>3 | Path:hsa04210 Apoptosis |
| 0.0022984 | 41 | 157 | 1.7579581 | Path:hsa04390 Hippo signaling pathway |

**Table S7: SILAC proteomics - PASEF acquisition table**

| #MS Type | Cycle Id | Start IM [1/K0] | End IM [1/K0] | Start Mass [m/z] | End Mass [m/z] | CE [eV] |
|---|---|---|---|---|---|---|
| MS1 | 0 | - | - | - | - | - |
| PASEF | 1 | 0.6 | 0.832 | 301.84 | 385.76 | - |
| PASEF | 1 | 0.832 | 1.5 | 621.8 | 631.29 | - |
| PASEF | 2 | 0.6 | 0.852 | 385.76 | 406.73 | - |
| PASEF | 2 | 0.852 | 1.5 | 631.29 | 640.98 | - |
| PASEF | 3 | 0.6 | 0.862 | 406.73 | 421.72 | - |
| PASEF | 3 | 0.862 | 1.5 | 640.98 | 650.82 | - |
| PASEF | 4 | 0.6 | 0.872 | 421.72 | 433.74 | - |
| PASEF | 4 | 0.872 | 1.5 | 650.82 | 660.83 | - |
| PASEF | 5 | 0.6 | 0.872 | 433.74 | 444.89 | - |
| PASEF | 5 | 0.872 | 1.5 | 660.83 | 670.84 | - |
| PASEF | 6 | 0.6 | 0.882 | 444.89 | 455.28 | - |
| PASEF | 6 | 0.882 | 1.5 | 670.84 | 681.02 | - |
| PASEF | 7 | 0.6 | 0.882 | 455.28 | 465.27 | - |
| PASEF | 7 | 0.882 | 1.5 | 681.02 | 691.88 | - |
| PASEF | 8 | 0.6 | 0.892 | 465.27 | 474.78 | - |
| PASEF | 8 | 0.892 | 1.5 | 691.88 | 703.33 | - |
| PASEF | 9 | 0.6 | 0.892 | 474.78 | 484.24 | - |
| PASEF | 9 | 0.892 | 1.5 | 703.33 | 715.38 | - |
| PASEF | 10 | 0.6 | 0.902 | 484.24 | 493.26 | - |
| PASEF | 10 | 0.902 | 1.5 | 715.38 | 727.37 | - |
| PASEF | 11 | 0.6 | 0.902 | 493.26 | 501.76 | - |
| PASEF | 11 | 0.902 | 1.5 | 727.37 | 740.4 | - |
| PASEF | 12 | 0.6 | 0.912 | 501.76 | 510.26 | - |
| PASEF | 12 | 0.912 | 1.5 | 740.4 | 754.03 | - |
| PASEF | 13 | 0.6 | 0.922 | 510.26 | 518.73 | - |
| PASEF | 13 | 0.922 | 1.5 | 754.03 | 768.36 | - |
| PASEF | 14 | 0.6 | 0.922 | 518.73 | 527.25 | - |
| PASEF | 14 | 0.922 | 1.5 | 768.36 | 782.91 | - |
| PASEF | 15 | 0.6 | 0.932 | 527.25 | 535.6 | - |
| PASEF | 15 | 0.932 | 1.5 | 782.91 | 798.44 | - |
| PASEF | 16 | 0.6 | 0.932 | 535.6 | 543.81 | - |
| PASEF | 16 | 0.932 | 1.5 | 798.44 | 815.91 | - |
| PASEF | 17 | 0.6 | 0.942 | 543.81 | 551.95 | - |
| PASEF | 17 | 0.942 | 1.5 | 815.91 | 833.42 | - |
| PASEF | 18 | 0.6 | 0.952 | 551.95 | 560.29 | - |
| PASEF | 18 | 0.952 | 1.5 | 833.42 | 853.44 | - |
| PASEF | 19 | 0.6 | 0.952 | 560.29 | 568.8 | - |
| PASEF | 19 | 0.952 | 1.5 | 853.44 | 875.73 | - |
| PASEF | 20 | 0.6 | 0.962 | 568.8 | 577.28 | - |

| | | | | | | |
|---|---|---|---|---|---|---|
| PASEF | 20 | 0.962 | 1.5 | 875.73 | 900.93 | - |
| PASEF | 21 | 0.6 | 0.972 | 577.28 | 585.83 | - |
| PASEF | 21 | 0.972 | 1.5 | 900.93 | 928.51 | - |
| PASEF | 22 | 0.6 | 0.982 | 585.83 | 594.33 | - |
| PASEF | 22 | 0.982 | 1.5 | 928.51 | 961.97 | - |
| PASEF | 23 | 0.6 | 0.992 | 594.33 | 603.3 | - |
| PASEF | 23 | 0.992 | 1.5 | 961.97 | 1004.99 | - |
| PASEF | 24 | 0.6 | 1.012 | 603.3 | 612.32 | - |
| PASEF | 24 | 1.012 | 1.5 | 1004.99 | 1067.01 | - |
| PASEF | 25 | 0.6 | 1.052 | 612.32 | 621.8 | - |
| PASEF | 25 | 1.052 | 1.5 | 1067.01 | 1199.55 | - |

**Table S8: SILAC proteomics - contrasts**

| RawfileName | SampleGroup |
|---|---|
| 251209ko02LM_lab_sample01_1_1_6945.d | GTD_WT_noTreat |
| 251209ko03LM_lab_sample02_1_1_6946.d | GTD_WT_noTreat |
| 251209ko04LM_lab_sample03_1_1_6947.d | GTD_WT_noTreat |
| 251209ko05LM_lab_sample04_1_1_6948.d | GTD_WT_IFNa |
| 251209ko06LM_lab_sample05_1_1_6949.d | GTD_WT_IFNa |
| 251209ko07LM_lab_sample06_1_1_6950.d | GTD_WT_IFNa |
| 251209ko14LM_lab_sample13_1_1_6957.d | GTD_BCPM_noTreat |
| 251209ko15LM_lab_sample14_1_1_6958.d | GTD_BCPM_noTreat |
| 251209ko16LM_lab_sample15_1_1_6959.d | GTD_BCPM_noTreat |
| 251209ko17LM_lab_sample16_1_1_6960.d | GTD_BCPM_IFNa |
| 251209ko18LM_lab_sample17_1_1_6961.d | GTD_BCPM_IFNa |
| 251209ko19LM_lab_sample18_1_1_6962.d | GTD_BCPM_IFNa |
| 251209ko26LM_lab_sample25_1_1_6969.d | GTD_PCM_noTreat |
| 251209ko27LM_lab_sample26_1_1_6970.d | GTD_PCM_noTreat |
| 251209ko28LM_lab_sample27_1_1_6971.d | GTD_PCM_noTreat |
| 251209ko29LM_lab_sample28_1_1_6972.d | GTD_PCM_IFNa |
| 251209ko30LM_lab_sample29_1_1_6973.d | GTD_PCM_IFNa |
| 251209ko31LM_lab_sample30_1_1_6974.d | GTD_PCM_IFNa |
| **Contrasts** | |
| GTD_WT_noTreat - GTD_WT_IFNa | - |
| GTD_PCM_noTreat - GTD_PCM_IFNa | - |
| GTD_BCPM_noTreat - GTD_BCPM_IFNa | - |
| GTD_WT_noTreat - GTD_PCM_noTreat | - |
| GTD_PCM_noTreat - GTD_BCPM_noTreat | - |
| GTD_WT_noTreat - GTD_BCPM_noTreat | - |
